## Supporting Information for "Structural insights into Cullin4-RING ubiquitin ligase remodelling by Vpr from simian immunodeficiency viruses"

Supporting Figures:

S1 Fig. Additional biochemical analysis of $Vpr_{mus}$-induced $CRL4^{DCAF1\text{-}CtD}$ specificity redirection towards SAMHD1.

S2 Fig. Components, controls and uncropped SDS-PAGE images of *in vitro* ubiquitylation reactions.

S3 Fig. Detailed crystal structure analysis of the DCAF1-CtD/$Vpr_{mus}$ complex.

S4 Fig. Cryo-EM analysis 1 of the $CRL4\text{-}NEDD8^{DCAF1\text{-}CtD}$/$Vpr_{mus}$/SAMHD1 complex.

S5 Fig. Cryo-EM analysis 2 and CLMS analysis of the $CRL4(\text{-}NEDD8)^{DCAF1\text{-}CtD}$/$Vpr_{mus}$/SAMHD1 complex.

S6 Fig. Multiple sequence alignment of Vpr/Vpx proteins, detailed structural comparison between $Vpr_{mus}$ and $Vpr_{HIV\text{-}1}$.

Supporting Tables:

S1 Table: X-ray data collection and refinement statistics.

S2 Table: Oligonucleotide primer sequences.

S3 Table: Expression constructs.

Supporting References

Supporting Files:

S1: PDB validation report 6zue

S2: PDB validation report 6zx9

S1 Fig

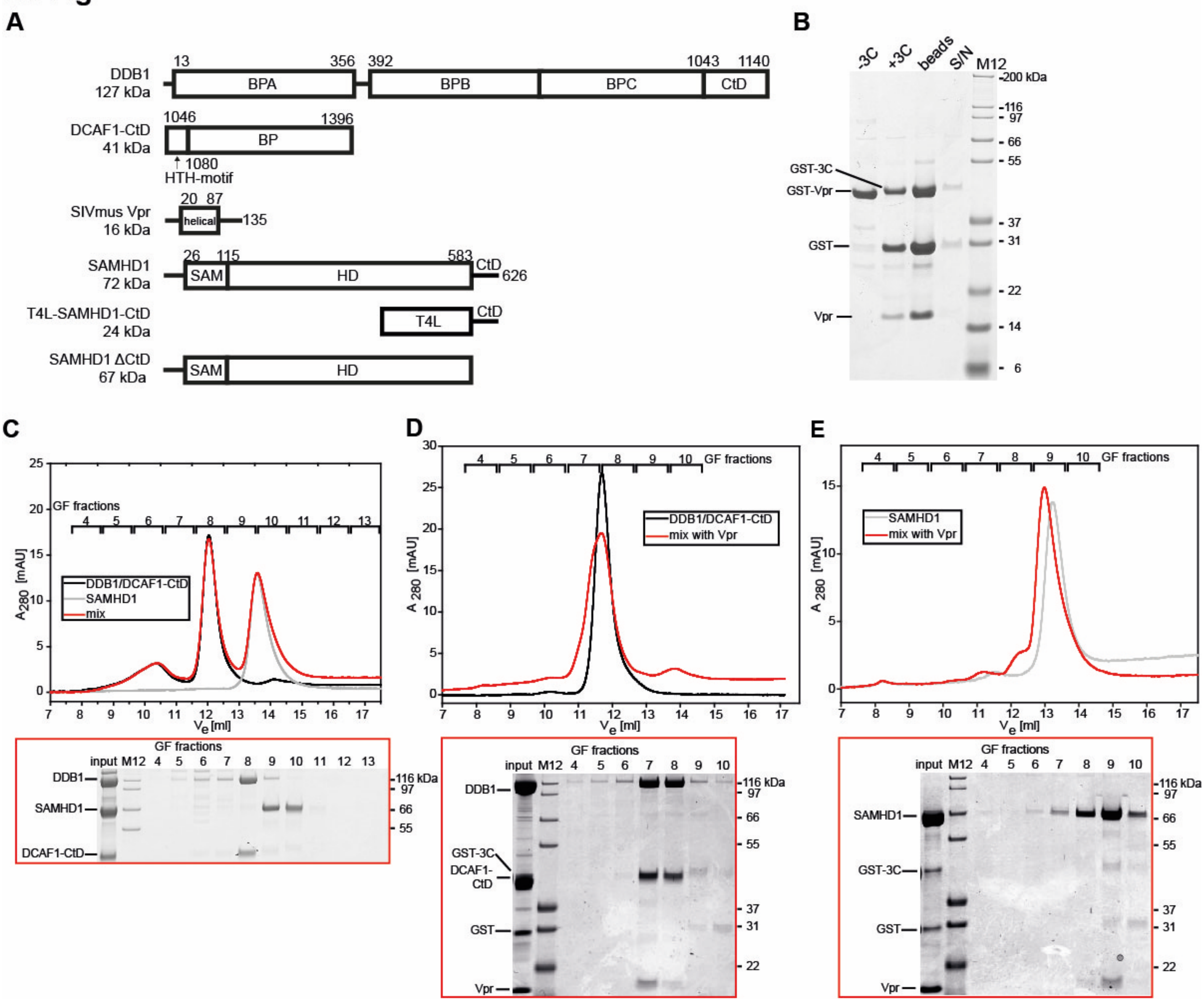

**S1 Fig. Additional biochemical analysis of $Vpr_{mus}$-induced $CRL4^{DCAF1\text{-}CtD}$ specificity redirection towards SAMHD1.**

(**A**) Schematic view of the protein constructs used in biochemical analyses. BP – β-propeller domain, HD – histidine-aspartate domain, HTH – helix-turn-helix motif, SAM – sterile alpha motif. (**B**) SDS-PAGE analysis of GST-$Vpr_{mus}$. After treatment with 3C protease to remove the GST-tag (+3C) and GSH-Sepharose pull down to remove protease and tag, no $Vpr_{mus}$ is present in the eluted fraction (S/N) indicating that it interacts non-specifically with the GSH-Sepharose beads and/or becomes insoluble after tag removal. (**C-E**) Analytical GF analysis of DDB1/DCAF1-CtD incubated with SAMHD1 (**C**), DDB1/DCAF1-CtD incubated with $Vpr_{mus}$ (**D**) and SAMHD1 incubated with $Vpr_{mus}$ (**E**). SDS-PAGE of the corresponding GF fractions is shown below each chromatogram.

#### S2 Fig

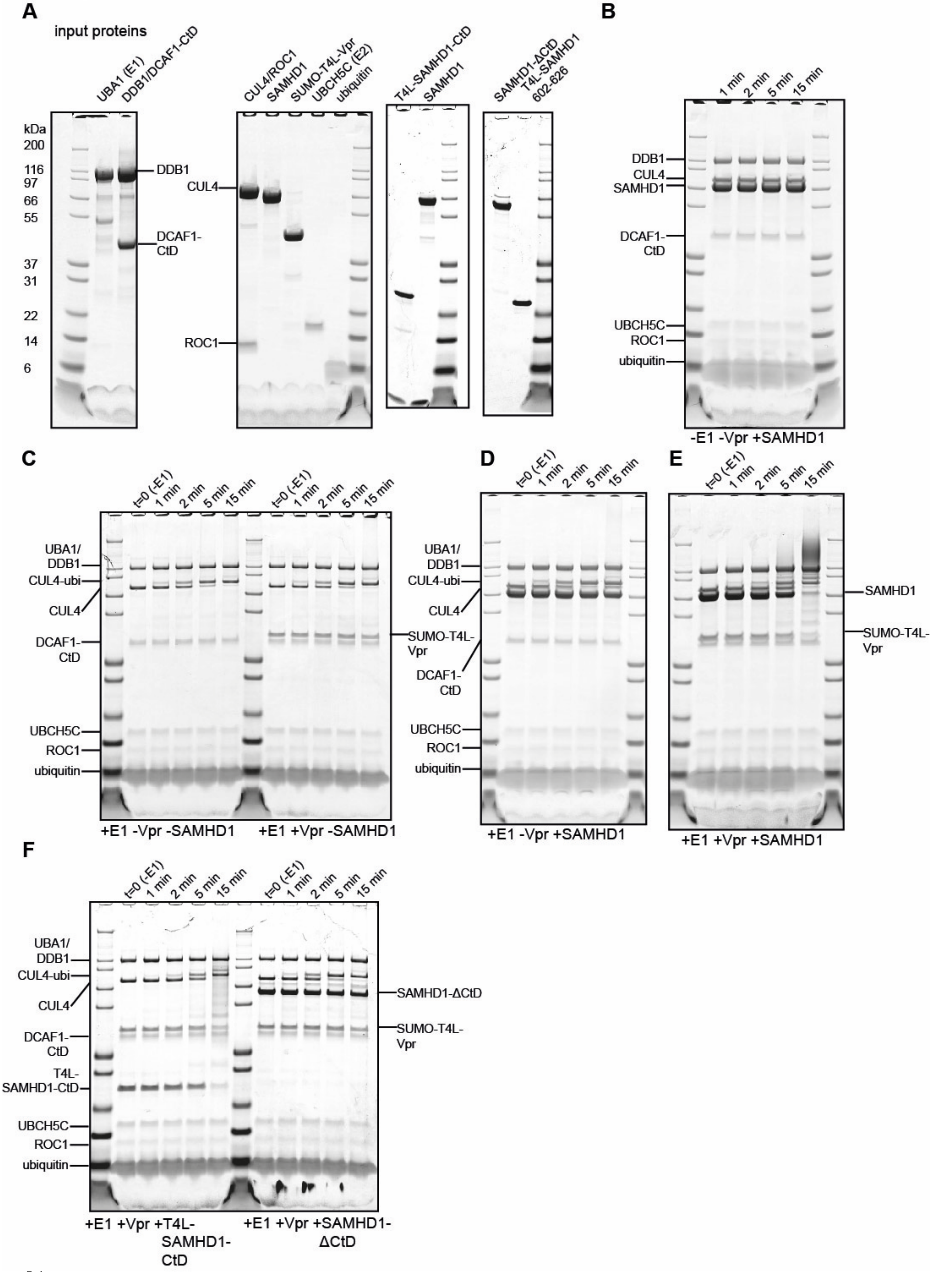
A
input proteins
UBA1 (E1)
DDB1/DCAF1-CtD
kDa
200
116
97
66
55
37
31
22
14
6
DDB1
DCAF1-CtD
CUL4/ROC1
SAMHD1
SUMO-T4L-Vpr
UBCH5C (E2)
ubiquitin
CUL4
ROC1
T4L-SAMHD1-CtD
SAMHD1
SAMHD1-ΔCtD
T4L-SAMHD1 602-626
B
1 min
2 min
5 min
15 min
DDB1
CUL4
SAMHD1
DCAF1-CtD
UBCH5C
ROC1
ubiquitin
-E1 -Vpr +SAMHD1
C
t=0 (-E1)
1 min
2 min
5 min
15 min
UBA1/DDB1
CUL4-ubi
CUL4
DCAF1-CtD
SUMO-T4L-Vpr
UBCH5C
ROC1
ubiquitin
+E1 -Vpr -SAMHD1
+E1 +Vpr -SAMHD1
D
t=0 (-E1)
1 min
2 min
5 min
15 min
UBA1/DDB1
CUL4-ubi
CUL4
DCAF1-CtD
UBCH5C
ROC1
ubiquitin
+E1 -Vpr +SAMHD1
E
t=0 (-E1)
1 min
2 min
5 min
15 min
SAMHD1
SUMO-T4L-Vpr
+E1 +Vpr +SAMHD1
F
t=0 (-E1)
1 min
2 min
5 min
15 min
UBA1/DDB1
CUL4-ubi
CUL4
DCAF1-CtD
T4L-SAMHD1-CtD
UBCH5C
ROC1
ubiquitin
SAMHD1-ΔCtD
SUMO-T4L-Vpr
+E1 +Vpr +T4L-SAMHD1-CtD
+E1 +Vpr +SAMHD1-ΔCtD

**S2 Fig. Components, controls and uncropped SDS-PAGE images of *in vitro* ubiquitylation reactions.**

(**A**) SDS-PAGE of individually purified protein components used in the *in vitro* ubiquitylation reactions. (**B, C**) Control reactions in the absence of indicated components. (**D-F**) Uncropped gels of reactions shown in Fig. 1C-F. All reactions were incubated at 37°C for the indicated times, stopped by addition of SDS sample buffer and separated on SDS-PAGE.

#### S3 Fig

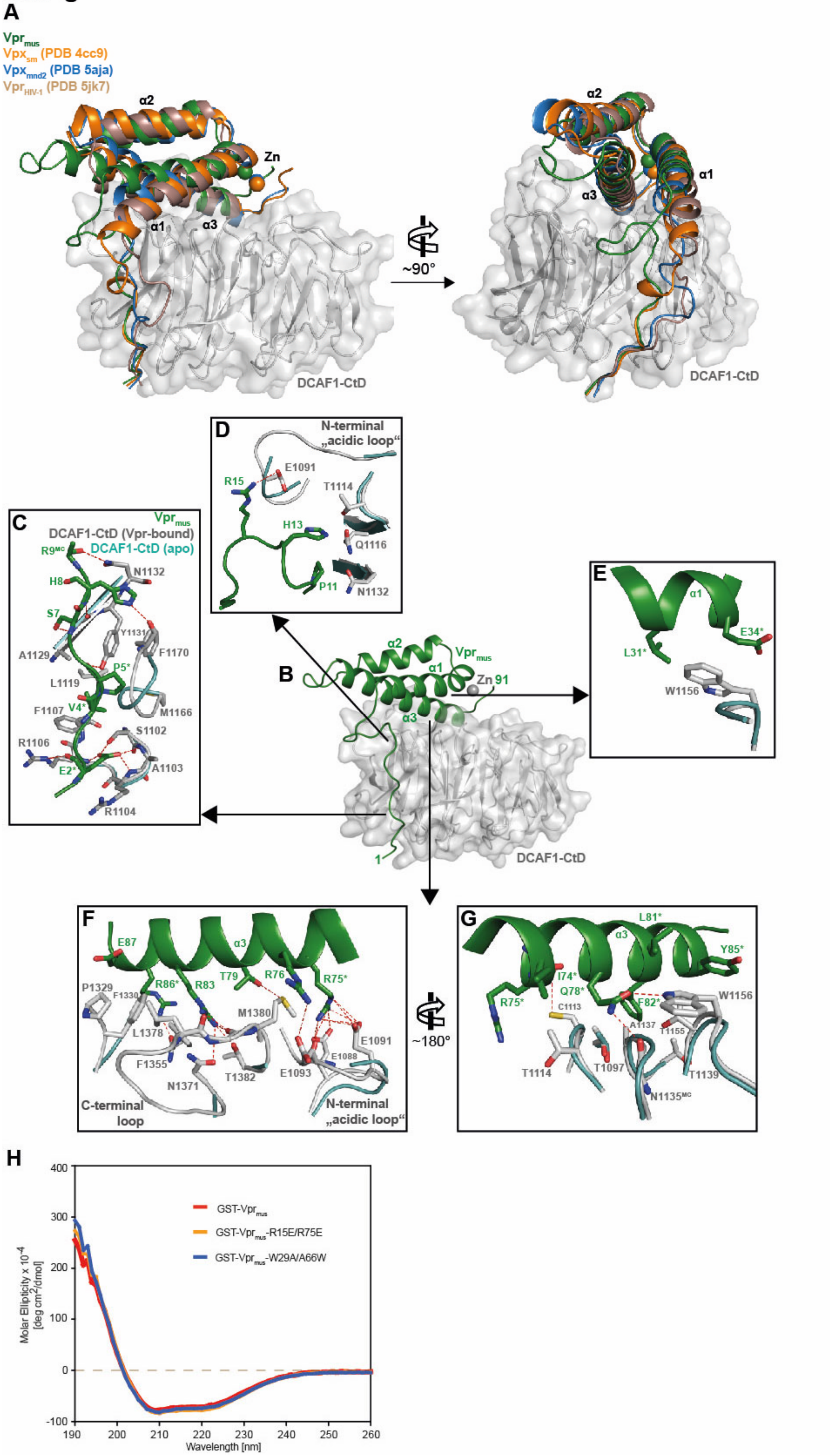
A
Vprmus
Vpxsm (PDB 4cc9)
Vpxmnd2 (PDB 5aja)
VprHIV-1 (PDB 5jk7)
α2
α1
α3
Zn
~90°
DCAF1-CtD
B
Vprmus
Zn 91
1
C
DCAF1-CtD (Vpr-bound)
DCAF1-CtD (apo)
D
N-terminal „acidic loop“
E
F
C-terminal loop
G
~180°
H
GST-Vprmus
GST-Vprmus-R15E/R75E
GST-Vprmus-W29A/A66W
Molar Ellipticity x 10^-4 [deg cm^2/dmol]
Wavelength [nm]

**S3 Fig. Detailed crystal structure analysis of the DDB1/DCAF1-CtD/$Vpr_{mus}$ complex.**

(**A**) Superposition of the $Vpr_{mus}$ (green cartoon)/DCAF1-CtD complex with $Vpx_{sm}$ (orange cartoon, PDB: 4cc9) [1], $Vpx_{mnd}$ (blue cartoon, PDB: 5aja) [2] and $Vpr_{HIV-1}$ (light brown cartoon, PDB: 5jk7) [3]. Structures have been aligned with respect to their DCAF1 BP domains but only the DCAF1-CtD from the $Vpr_{mus}$ complex is shown for clarity (grey cartoon and semi-transparent surface). (**B-G**) Details of the DCAF1-CtD/$Vpr_{mus}$ interaction. (**B**) The structure of the complex is shown in the same orientation as Fig. 2A, left panel. The insets (**C**-**G**) show individual interaction areas in more detail, $Vpr_{mus}$ (green), $Vpr_{mus}$-bound DCAF1-CtD (grey) and apo-DCAF1-CtD (light blue). Selected amino acid residues, that make intermolecular interactions, are shown as sticks, and hydrogen bonds/electrostatic interactions as dashed red lines. $Vpr_{mus}$ residues with asterisks are type-conserved within all Vpr/Vpx proteins. (**H**) Circular dichroism (CD) spectra of GST-$Vpr_{mus}$ and GST-$Vpr_{mus}$ variants R15E/R75E and W29A/A66W.

#### S4 Fig

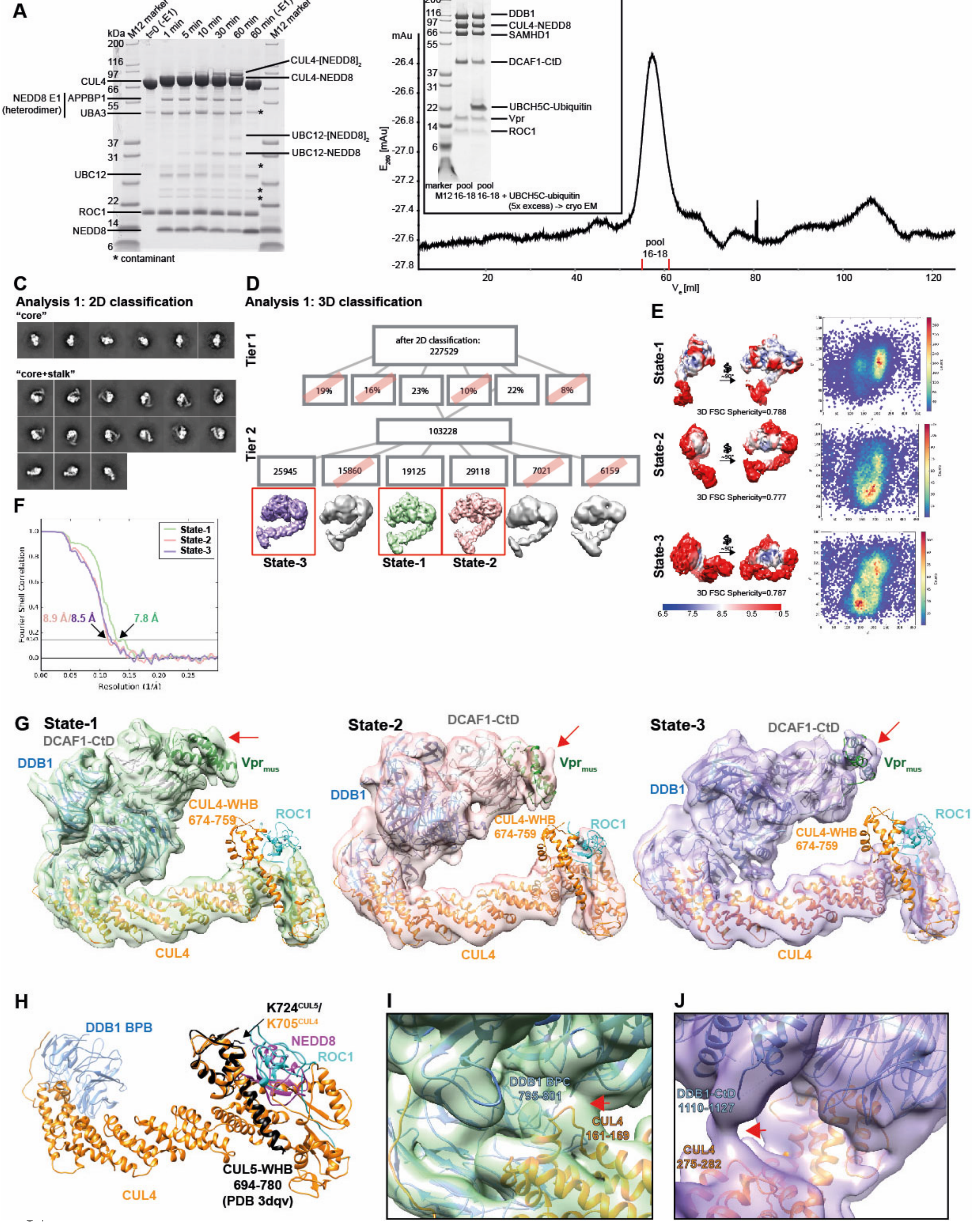
A
kDa
M12 marker
t=0 (-E1)
1 min
5 min
10 min
30 min
60 min
60 min (-E1)
M12 marker
CUL4-[NEDD8]2
CUL4-NEDD8
CUL4
NEDD8 E1
(heterodimer)
APPBP1
UBA3
UBC12-[NEDD8]2
UBC12-NEDD8
UBC12
ROC1
NEDD8
* contaminant
B
marker M12
pool 16-18
pool 16-18 + UBCH5C-ubiquitin (5x excess) -> cryo EM
DDB1
CUL4-NEDD8
SAMHD1
DCAF1-CtD
UBCH5C-Ubiquitin
Vpr
ROC1
mAu
E280 [mAu]
pool 16-18
Ve [ml]
C
Analysis 1: 2D classification
"core"
"core+stalk"
D
Analysis 1: 3D classification
Tier 1
after 2D classification: 227529
19%
16%
23%
10%
22%
8%
Tier 2
103228
25945
15860
19125
29118
7021
6159
State-3
State-1
State-2
E
State-1
3D FSC Sphericity=0.788
State-2
3D FSC Sphericity=0.777
State-3
3D FSC Sphericity=0.787
F
State-1
State-2
State-3
Fourier Shell Correlation
8.9 Å/8.5 Å
7.8 Å
Resolution (1/Å)
G
State-1
DCAF1-CtD
DDB1
Vprmus
CUL4-WHB 674-759
ROC1
CUL4
State-2
State-3
H
DDB1 BPB
K724CUL5/
K705CUL4
NEDD8
ROC1
CUL5-WHB 694-780 (PDB 3dqv)
CUL4
I
DDB1 BPC 795-801
CUL4 161-169
J
DDB1-CtD 1110-1127
CUL4 275-282

**S4 Fig. Cryo-EM analysis 1 of the CRL4-NEDD8$^{DCAF1\text{-}CtD}$/Vpr$_{mus}$/SAMHD1 complex.**

(**A**) *In vitro* neddylation of CUL4/ROC1. Protein was mixed with purified neddylation-E1 (APPBP1/UBA3 heterodimer), E2 (UBC12) and NEDD8. The reaction was incubated at 25°C, samples were taken at indicated times, stopped by addition of SDS sample buffer and separated on SDS-PAGE. (**B**) GF analysis of the CUL4-NEDD8/ROC1/DDB1/DCAF1-CtD/Vpr$_{mus}$/SAMHD1 complex with pooled fractions indicated. A 5x molar excess of UBCH5C-ubiquitin was added before plunge-freezing for cryo-EM experiments, in an attempt to stabilise the assembly. However, no density in any of the reconstructions could be assigned to UBCH5C-ubiquitin, indicating low binding affinity and/or heterogeneity in its mode of binding. (**C**) 2D class averages depicting either "core" or "core+stalk" classes of analysis 2. (**D**) 3D sorting tree after 2D classification. Conformational states-1, -2 and -3 are indicated. (**E**) Local resolution and Euler distribution of states-1, -2 and -3. (**F**) FSC curves for state-1, -2 and -3 reconstructions. (**G**) Side-by-side comparison of state-1, -2 and -3 reconstructions, coloured as in Fig. 3. Molecular models of the DDB1/DCAF1-CtD/Vpr$_{mus}$ crystal structure and CUL4/ROC1 (PDB 2hye) [4] have been fitted as rigid bodies into the volumes and are shown as cartoons. DDB1/DCAF1-CtD/Vpr$_{mus}$ is coloured as in Fig. 4, CUL4 is coloured yellow and ROC1 cyan. All states show additional density corresponding to SAMHD1-CtD, indicated by the red arrows. (**H**) Superposition of the neddylated CUL5 C-terminal WHB domain (black cartoon, PDB 3dqv) [5] on the CUL4 WHB (PDB 2hye), coloured as in **A**. Respective lysine residues, which are covalently modified with NEDD8, are indicated. (**I**, **J**) Detailed view of state-1 (I) and state-3 (J) cryo-EM density. Red arrows indicate contacts between CUL4A (orange cartoon) and DDB1 BPA/BPC/CtD (blue cartoon).

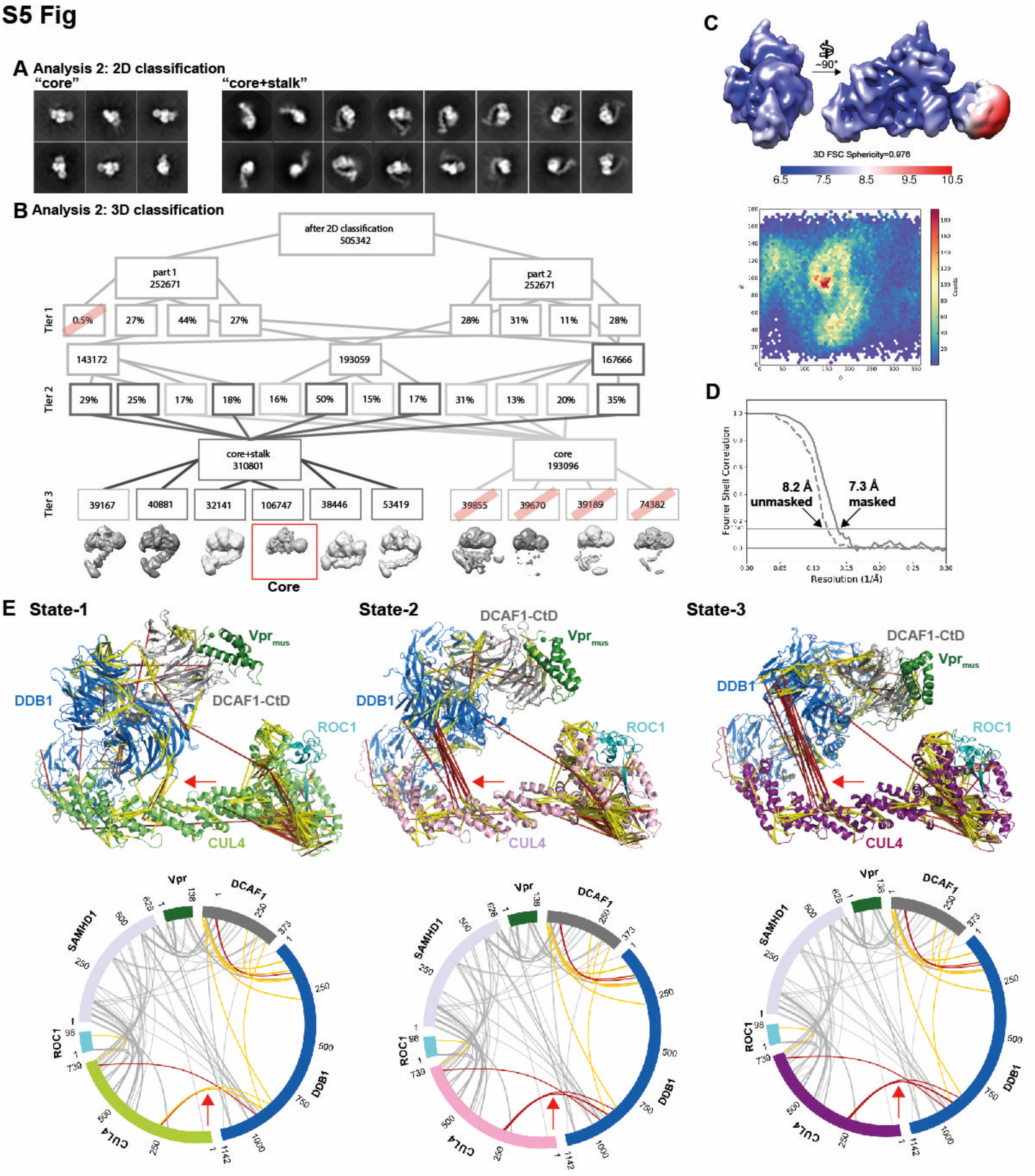

76

77 **S5 Fig. Cryo-EM analysis 2 and CLMS analysis of the CRL4(-NEDD8)$^{DCAF1\text{-}CtD}$/ Vpr$_{mus}$ /SAMHD1**

78 **complex.**

(**A**) 2D class averages depicting either CRL4-NEDD8$^{DCAF1\text{-}CtD}$/Vpr$_{mus}$/SAMHD1 “core” or “core+stalk” classes of analysis 1. (**B**) Sorting tree after 2D classification. In Tier 3, the core reconstruction was identified, containing 106,747 particle images (red box). (**C**) Local resolution of the core reconstruction after refinement, indicating a resolution range from 6.5 Å in the hydrophobic interior of DDB1 to 10.5Å in the DDB1 BPB domain. Below, the Euler distribution is shown. (**D**) FSC curve of the core reconstruction after refinement. (**E**) Upper panel: CRL4$^{DCAF1\text{-}CtD}$/Vpr$_{mus}$/SAMHD1 cross-links, identified by CLMS, mapped on molecular models representing state-1, -2 and -3. Satisfied crosslinks (<25 Å) are coloured yellow, violated crosslinks red. Red arrows indicate a subset of cross-links between DDB1 and CUL4, whose distance restraints are satisfied in state-1, and increasingly violated in states-2 and -3. Lower panel: circle plot of CLMS data for states-1, -2 and -3, using the same colour scheme as in the upper panel. Grey lines represent crosslinks between residues that are not present in the molecular models. Only crosslinks between subunits are displayed.

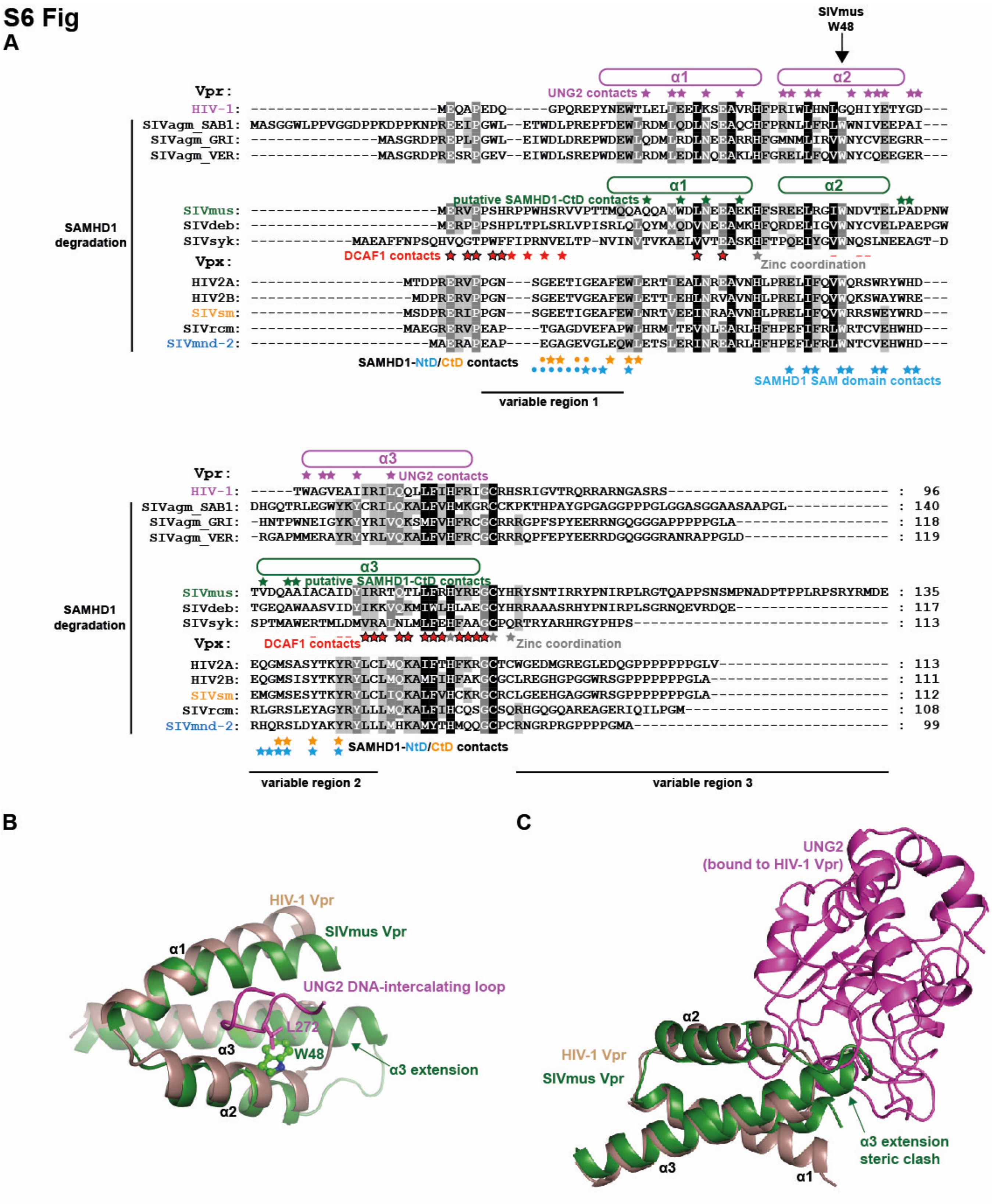

**S6 Fig. Multiple sequence alignment of Vpr/Vpx proteins, detailed structural comparison between $Vpr_{mus}$ and $Vpr_{HIV\text{-}1}$.**

(**A**) Sequence alignment of indicated Vpr and Vpx proteins. Helices are indicated by the boxes above the amino acid sequences for $Vpr_{HIV-1}$ (pink) and $Vpr_{mus}$ (green). $Vpr_{HIV-1}$ side chains involved in UNG2-binding are indicated with pink asterisks. $Vpr_{mus}$ side chains putatively involved in SAMHD1-CtD-binding are indicated by green asterisks. $Vpx_{sm}$ side chains targeting SAMHD1-CtD are indicated with orange asterisks, and $Vpx_{mnd2}$ side chains contacting N-terminal SAMHD1 domains are highlighted with blue asterisks. Red symbols mark $Vpr_{mus}$ side chains involved in DCAF1-CtD-binding. The non-outlined symbols indicate DCAF1-CtD-contacting side chains unique to $Vpr_{mus}$, and dashes show DCAF1-binding side chains, which are in contact with DCAF1-CtD in other Vpr/Vpx structures, but not in $Vpr_{mus}$. Grey asterisks mark Vpr/Vpx side chains involved in zinc coordination. (**B**) Structural alignment of $Vpr_{HIV-1}$ (PDB 5jk7 [3], light brown) in complex with UNG2 and $Vpr_{mus}$ (green). Protein backbone is shown in cartoon representation. For clarity, only the DNA-intercalating loop of UNG2 is shown (pink), which inserts into a hydrophobic pocket created by the $Vpr_{HIV-1}$ helix bundle. Note the steric clash between UNG2 side chain L272 and $Vpr_{mus}$ residue W48 in the structural superposition. (**C**) Alternative view of the structural alignment of $Vpr_{HIV-1}$ (light brown) in complex with UNG2 (pink) and $Vpr_{mus}$ (green). Note the steric clash between UNG2 and the extended Helix-3 of $Vpr_{mus}$.

**S1 Table: X-ray data collection and refinement statistics.**

| Sample | DDB1/DCAF1-CtD | DDB1/DCAF1-CtD/T4L-$Vpr_{mus}$ 1-92 | |
|---|---|---|---|
| PDB code | 6zue | - | 6zx9 |
| *Data collection* | | | |
| Space group | I222 | $P2_12_12$ | $P2_12_12$ |
| Cell dimensions | | | |
| a, b, c (Å) | 117.38, 153.63, 223.16 | 266.91, 95.94, 99.35 | 265.90, 95.54, 98.35 |
| α, β, γ (°) | 90, 90, 90 | 90, 90, 90 | 90, 90, 90 |
| Resolution range (Å) | 50.00 (3.28)* – 3.09 | 50.00 (3.83) – 3.61 | 79.07 (2.56) – 2.52 |
| $R_{merge}$ (%) | 8.8 (120.0) | 32.9 (136.1) | 9.9 (162.6) |
| $CC_{1/2}$** | 99.9 (85.3) | 100.0 (35.4) | 99.6 (48.0) |
| I/σ(I) | 13.0 (1.4) | 5.9 (1.3) | 9.8 (1.1) |
| Completeness (%) | 96.8 (97.1) | 99.3 (96.2) | 99.9 (99.8) |
| Redundancy | 4.8 (4.9) | 6.4 (6.1) | 6.6 (6.7) |
| *Refinement* | | | |
| Resolution range (Å) | 48.60 (3.18) – 3.09 | - | 79.07 (2.55) – 2.52 |
| No. reflections | 35922 (2815) | - | 84808 (2880) |
| $R_{work}/R_{free}$ (%) | 22.0/27.9 (35.9/43.5) | - | 21.6/26.1 (41.1/44.6) |
| No. atoms | | | |
| Protein | 11279 | - | 13556 |
| Ligand/ion | - | - | 49 |
| Water | 10 | - | 253 |
| B-factors | | | |
| Protein | 114.6 | - | 75.5 |
| Ligand/ion | - | - | 77.2 |
| Water | 71.7 | - | 54.9 |
| R.m.s. deviations | | | |
| Bond lengths (Å) | 0.004 | - | 0.004 |
| Bond angles (°) | 0.759 | - | 0.794 |

*Numbers in parentheses account for the high-resolution shell

**defined in [6]

**S2 Table: Oligonucleotide primer sequences.**

| Restriction enzyme, insert | Sequence | Destination plasmid (restriction enzymes) |
|---|---|---|
| NcoI hsDCAF1 1046 fw. | ggcCCATGGCGccaataaactttacgtcaaggc | pTriEx-6 (NcoI/SacI) |
| SacI hsDCAF1 1396 rev. | ggcGAGCTCctctgccagacgctgcctgcc | |
| XmaI T4L (E11H) 2 fw. | ggcCCCGGGaacatttttgaaatgctgcgtattgatg | pHisSUMO (XmaI/NotI) |
| NotI T4L (E11H) 164 rev. | ggcGCGGCCGCcaggtttttataggcatcccatgtg | |
| AgeI rhSAMHD1 1 fw. | atattACCGGTatgcagcaagccgactcc | pHisSUMO (XmaI/NotII) |
| *NotI rhSAMHD1 583 rev.* | taattGCGGCCGCTTAatcctgaggcttggtgaaatttc | |
| *NotI rhSAMHD1 626 rev.* | taattGCGGCCGCTTActttgggtcatctttaaaaagc | |
| NotI rhSAMHD1 582 fw. | ggcGCGGCCGCAcaggatggtgatgttattgcacc | pHisSUMO-T4L (E11H) (NotI/SacI) |
| *SacI rhSAMHD1 626 rev.* | ggcGAGCTCTTAttatttcggatcatctttaaacagctg | |
| *XmaI Vpr$_{mus}$ 1 fw.* | ggcCCCGGGatggaacgtgttccgcctagcc | pET49b (XmaI/XhoI) |
| *XhoI Vpr$_{mus}$ 135 rev.* | ggcCTCGAGTTAttattcatccatacgataacggctc | |
| *NotI Vpr$_{mus}$ 1 fw.* | ggcGCGGCCGCAatggaacgtgttccgcctagcc | pHisSUMO-T4L (E11H) (NotI/SacI) |
| *SacI Vpr$_{mus}$ 92 rev.* | ggcGAGCTCTTATTAgcggtgataacaaccttcacgataatg | |
| *Vpr$_{mus}$ R15E fw.* | GGCATAGCGAAGTTGTTCCGACCACC | pET49b-*Vpr$_{mus}$* |
| *Vpr$_{mus}$ R15E rev.* | GGTCGGAACAACTTGGCTATGCCAAGG | |
| *Vpr$_{mus}$ R75E fw.* | GATTATATTGAACGTACCCAGACCCTGCTG | |
| *Vpr$_{mus}$ R75E rev.* | GTCTGGGTACGTTCAATATAATCAATGGCAC | |
| *Vpr$_{mus}$ W29A fw.* | GCACAGCAGGCCATGGCGGATCTGAATGAAGAAGCA | |
| *Vpr$_{mus}$ W29A rev.* | TTCTTCATTCAGATCCGCCATGGCCTGCTGTGCCTG | |
| *Vpr$_{mus}$ A66W fw.* | GGACCGTTGATCAGGCATGGATTGCATGTGCCATTGATTATATTC | |
| *Vpr$_{mus}$ A66W rev.* | CAATGGCACATGCAATCCATGCCTGATCAACGGTCCAATTC | |
| *NdeI ROC1 1 fw.* | ggcCATATGgcggcagcgatggatgtgg | pRSF-Duet-1 (NdeI/XhoI) |
| *XhoI ROC1 108 rev.* | ggcCTCGAGCTActagtgcccatacttttggaattc | |
| *BamHI hsCUL4A 2 fw.* | ggcGGATCCGgcggacgaggccccgcgg | pRSF-Duet-1-ROC1 (12-108) (BamHI/NotI) |
| *NotI hsCUL4A 759 rev.* | ggcGCGGCCGCTCAtcaggccacgtagtggtactgattc | |
| *BamHI UBCH5C 1 fw.* | ggcGGATCCatggcgctgaaacggattaataag | pGex6P1 (BamHI/NotI) |
| *NotI UBCH5C 147 rev.* | ggcGCGGCCGCTCAcatggcatacttctgagtcc | |

**S3 Table: Expression constructs.**

| Protein | UniProt ID | Vector | Expression system |
|---|---|---|---|
| Homo sapiens (hs) DDB1<br>(full length (fl)) | Q16531 | pAcGHLT-B | Sf9 |
| hsDCAF1-CtD<br>(residues 1046-1396) | Q9Y4B6 | pTri-Ex-6 | Sf9 |
| *Macaca mulatta*<br>*(Rhesus macaque, rh)* SAMHD1<br>(fl)<br>(ΔCtD, residues 1-583) | G7N4W9 | pHisSUMO | E. Coli Rosetta 2 (DE3) |
| T4L(variant E11H)-rhSAMHD1-CtD<br>(residues 582-626) | G7N4W9 | pHisSUMO | E. Coli Rosetta 2 (DE3) |
| *SIVmus* Vpr (WT and variants R15E/R75E; W29A/A66W)<br>(fl) | A4UDG5 | pET49b | E. Coli Rosetta 2 (DE3) |
| T4L(variant E11H)-*SIVmus* Vpr (residues 1-92) | A4UDG5 | pHisSUMO | E. Coli Rosetta 2 (DE3) |
| hs*Cullin4A (CUL4A) (residues 38-759)*<br>*hsROC1 (residues 12-108 )* | CUL4A: Q13619<br>ROC1: P62877 | pRSF-Duet-1 (Co-expression) | E. Coli Rosetta 2 (DE3) |
| *Mus musculus (mm)*<br>*mmUBA1*<br>(fl) | Q02053 | pET28 | E. Coli Rosetta 2 (DE3) |
| *hsUBCH5C*<br>*(fl)* | P61077 | pGex6P1 | E. Coli Rosetta 2 (DE3) |
| *hsUBA3*<br>*(fl)*<br>*hsAPPBP1*<br>*(fl)* | UBA3: Q8TBC4-2<br>APPBP1: Q13564 | pOPC (Co-expression) | E. Coli Rosetta 2 (DE3) |
| *hsUBC12*<br>*(fl)* | P61081 | pGex6P2 | E. Coli Rosetta 2 (DE3) |

### Full wwPDB X-ray Structure Validation Report (i)

Jul 23, 2020 – 11:46 AM BST

| | | |
|---|---|---|
| PDB ID | : | 6ZUE |
| Title | : | Crystal structure of human DDB1 bound to human DCAF1 (amino acid residues 1046-1396) |
| Deposited on | : | 2020-07-22 |
| Resolution | : | 3.09 Å(reported) |

This is a Full wwPDB X-ray Structure Validation Report.

This report is produced by the wwPDB biocuration pipeline after annotation of the structure.

We welcome your comments at **
A user guide is available at
https://www.wwpdb.org/validation/2017/XrayValidationReportHelp
with specific help available everywhere you see the (i) symbol.

CONFIDENTIAL VALIDATION REPORT

The following versions of software and data (see references (i)) were used in the production of this report:

| | | |
|---|---|---|
| MolProbity | : | 4.02b-467 |
| Xtriage (Phenix) | : | 1.13 |
| EDS | : | 2.13 |
| Percentile statistics | : | 20191225.v01 (using entries in the PDB archive December 25th 2019) |
| Refmac | : | 5.8.0158 |
| CCP4 | : | 7.0.044 (Gargrove) |
| Ideal geometry (proteins) | : | Engh & Huber (2001) |
| Ideal geometry (DNA, RNA) | : | Parkinson et al. (1996) |
| Validation Pipeline (wwPDB-VP) | : | 2.13 |

### 1 Overall quality at a glance (i)

The following experimental techniques were used to determine the structure:
*X-RAY DIFFRACTION*

The reported resolution of this entry is 3.09 Å.

Percentile scores (ranging between 0-100) for global validation metrics of the entry are shown in the following graphic. The table shows the number of entries on which the scores are based.

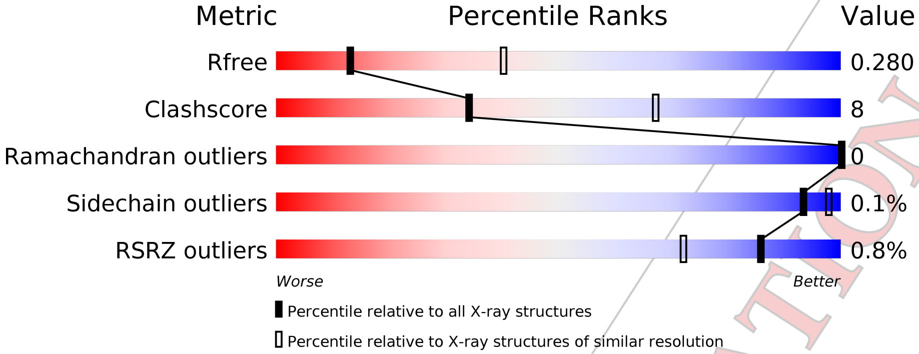

| Metric | Whole archive (#Entries) | Similar resolution (#Entries, resolution range(Å)) |
|---|---|---|
| $R_{free}$ | 130704 | 1094 (3.10-3.10) |
| Clashscore | 141614 | 1184 (3.10-3.10) |
| Ramachandran outliers | 138981 | 1141 (3.10-3.10) |
| Sidechain outliers | 138945 | 1141 (3.10-3.10) |
| RSRZ outliers | 127900 | 1067 (3.10-3.10) |

The table below summarises the geometric issues observed across the polymeric chains and their fit to the electron density. The red, orange, yellow and green segments on the lower bar indicate the fraction of residues that contain outliers for >=3, 2, 1 and 0 types of geometric quality criteria respectively. A grey segment represents the fraction of residues that are not modelled. The numeric value for each fraction is indicated below the corresponding segment, with a dot representing fractions <=5% The upper red bar (where present) indicates the fraction of residues that have poor fit to the electron density. The numeric value is given above the bar.

| Mol | Chain | Length | Quality of chain |
|---|---|---|---|
| 1 | A | 1142 | 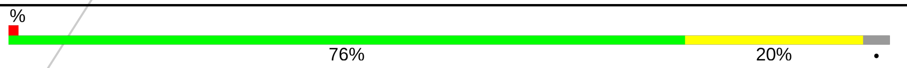 |
| 2 | B | 373 | 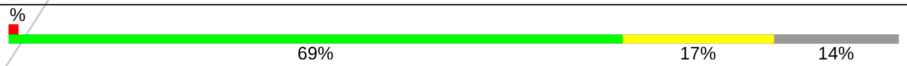 |

CONFIDENTIAL VALIDATION REPORT

### 2 Entry composition (i)

There are 3 unique types of molecules in this entry. The entry contains 11289 atoms, of which 0 are hydrogens and 0 are deuteriums.

In the tables below, the ZeroOcc column contains the number of atoms modelled with zero occupancy, the AltConf column contains the number of residues with at least one atom in alternate conformation and the Trace column contains the number of residues modelled with at most 2 atoms.

- Molecule 1 is a protein called DNA damage-binding protein 1.

| Mol | Chain | Residues | Atoms | | | | | ZeroOcc | AltConf | Trace |
|---|---|---|---|---|---|---|---|---|---|---|
| 1 | A | 1106 | Total<br>8689 | C<br>5514 | N<br>1462 | O<br>1665 | S<br>48 | 0 | 1 | 0 |

There are 2 discrepancies between the modelled and reference sequences:

| Chain | Residue | Modelled | Actual | Comment | Reference |
|---|---|---|---|---|---|
| A | -1 | GLY | - | expression tag | UNP Q16531 |
| A | 0 | SER | - | expression tag | UNP Q16531 |

- Molecule 2 is a protein called DDB1- and CUL4-associated factor 1.

| Mol | Chain | Residues | Atoms | | | | | ZeroOcc | AltConf | Trace |
|---|---|---|---|---|---|---|---|---|---|---|
| 2 | B | 322 | Total<br>2590 | C<br>1642 | N<br>455 | O<br>477 | S<br>16 | 0 | 1 | 0 |

There are 22 discrepancies between the modelled and reference sequences:

| Chain | Residue | Modelled | Actual | Comment | Reference |
|---|---|---|---|---|---|
| B | 1045 | MET | - | initiating methionine | UNP Q9Y4B6 |
| B | 1397 | GLU | - | expression tag | UNP Q9Y4B6 |
| B | 1398 | LEU | - | expression tag | UNP Q9Y4B6 |
| B | 1399 | ALA | - | expression tag | UNP Q9Y4B6 |
| B | 1400 | LEU | - | expression tag | UNP Q9Y4B6 |
| B | 1401 | VAL | - | expression tag | UNP Q9Y4B6 |
| B | 1402 | PRO | - | expression tag | UNP Q9Y4B6 |
| B | 1403 | ARG | - | expression tag | UNP Q9Y4B6 |
| B | 1404 | GLY | - | expression tag | UNP Q9Y4B6 |
| B | 1405 | SER | - | expression tag | UNP Q9Y4B6 |
| B | 1406 | SER | - | expression tag | UNP Q9Y4B6 |
| B | 1407 | ALA | - | expression tag | UNP Q9Y4B6 |
| B | 1408 | HIS | - | expression tag | UNP Q9Y4B6 |
| B | 1409 | HIS | - | expression tag | UNP Q9Y4B6 |

*Continued on next page...*

CONFIDENTIAL VALIDATION REPORT

*Continued from previous page...*

| Chain | Residue | Modelled | Actual | Comment | Reference |
|---|---|---|---|---|---|
| B | 1410 | HIS | - | expression tag | UNP Q9Y4B6 |
| B | 1411 | HIS | - | expression tag | UNP Q9Y4B6 |
| B | 1412 | HIS | - | expression tag | UNP Q9Y4B6 |
| B | 1413 | HIS | - | expression tag | UNP Q9Y4B6 |
| B | 1414 | HIS | - | expression tag | UNP Q9Y4B6 |
| B | 1415 | HIS | - | expression tag | UNP Q9Y4B6 |
| B | 1416 | HIS | - | expression tag | UNP Q9Y4B6 |
| B | 1417 | HIS | - | expression tag | UNP Q9Y4B6 |

- Molecule 3 is water.

| Mol | Chain | Residues | Atoms | ZeroOcc | AltConf |
|---|---|---|---|---|---|
| 3 | A | 9 | Total O<br>9 9 | 0 | 0 |
| 3 | B | 1 | Total O<br>1 1 | 0 | 0 |

CONFIDENTIAL VALIDATION REPORT

### 3 Residue-property plots (i)

These plots are drawn for all protein, RNA, DNA and oligosaccharide chains in the entry. The first graphic for a chain summarises the proportions of the various outlier classes displayed in the second graphic. The second graphic shows the sequence view annotated by issues in geometry and electron density. Residues are color-coded according to the number of geometric quality criteria for which they contain at least one outlier: green = 0, yellow = 1, orange = 2 and red = 3 or more. A red dot above a residue indicates a poor fit to the electron density (RSRZ > 2). Stretches of 2 or more consecutive residues without any outlier are shown as a green connector. Residues present in the sample, but not in the model, are shown in grey.

- Molecule 1: DNA damage-binding protein 1

Chain A:

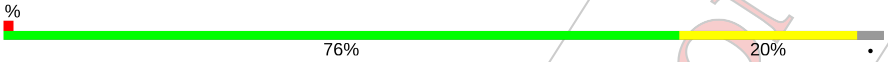

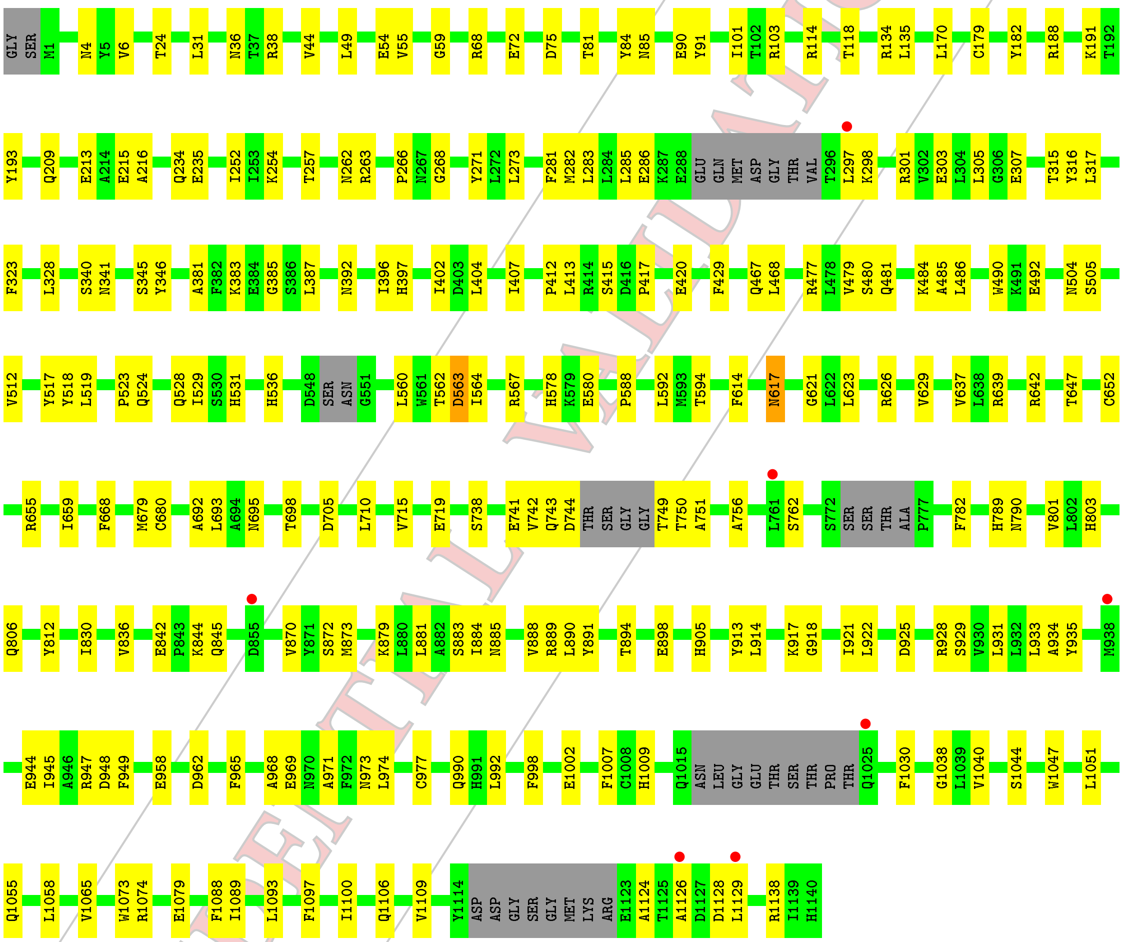

- Molecule 2: DDB1- and CUL4-associated factor 1

Chain B:

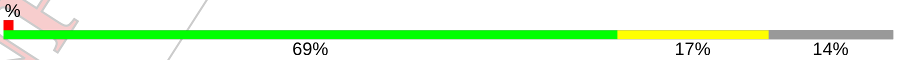

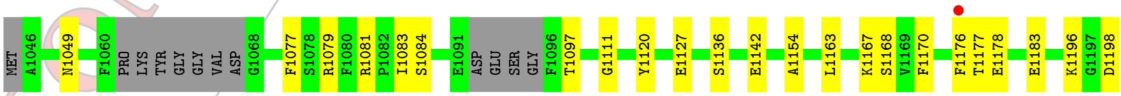

CONFIDENTIAL VALIDATION REPORT

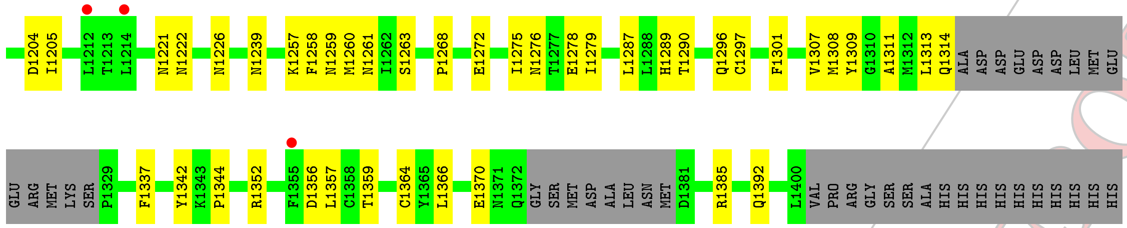

CONFIDENTIAL VALIDATION REPORT

### 4 Data and refinement statistics (i)

| Property | Value | Source |
|---|---|---|
| Space group | I 2 2 2 | Depositor |
| Cell constants<br>a, b, c, $\alpha$, $\beta$, $\gamma$ | 117.38Å 153.63Å 223.16Å<br>90.00° 90.00° 90.00° | Depositor |
| Resolution (Å) | 48.63 – 3.09<br>48.63 – 3.09 | Depositor<br>EDS |
| % Data completeness<br>(in resolution range) | 96.6 (48.63-3.09)<br>96.8 (48.63-3.09) | Depositor<br>EDS |
| $R_{merge}$ | (Not available) | Depositor |
| $R_{sym}$ | (Not available) | Depositor |
| $< I/\sigma(I) >$ [1] | 0.97 (at 3.07Å) | Xtriage |
| Refinement program | PHENIX 1.12_2829 | Depositor |
| R, $R_{free}$ | 0.222 , 0.279<br>0.222 , 0.280 | Depositor<br>DCC |
| $R_{free}$ test set | 1786 reflections (4.96%) | wwPDB-VP |
| Wilson B-factor (Å$^2$) | 101.0 | Xtriage |
| Anisotropy | 0.593 | Xtriage |
| Bulk solvent $k_{sol}$(e/Å$^3$), $B_{sol}$(Å$^2$) | 0.30 , 75.5 | EDS |
| L-test for twinning[2] | $< \|L\| > = 0.48$, $< L^2 > = 0.31$ | Xtriage |
| Estimated twinning fraction | No twinning to report. | Xtriage |
| $F_o$,$F_c$ correlation | 0.95 | EDS |
| Total number of atoms | 11289 | wwPDB-VP |
| Average B, all atoms (Å$^2$) | 113.0 | wwPDB-VP |

Xtriage's analysis on translational NCS is as follows: *The largest off-origin peak in the Patterson function is 2.74% of the height of the origin peak. No significant pseudotranslation is detected.*

[1] Intensities estimated from amplitudes.

[2] Theoretical values of $< |L| >$, $< L^2 >$ for acentric reflections are 0.5, 0.333 respectively for untwinned datasets, and 0.375, 0.2 for perfectly twinned datasets.

CONFIDENTIAL VALIDATION REPORT

### 5 Model quality (i)

#### 5.1 Standard geometry (i)

The Z score for a bond length (or angle) is the number of standard deviations the observed value is removed from the expected value. A bond length (or angle) with $|Z| > 5$ is considered an outlier worth inspection. RMSZ is the root-mean-square of all Z scores of the bond lengths (or angles).

| Mol | Chain | Bond lengths | | Bond angles | |
|---|---|---|---|---|---|
| | | RMSZ | #\|Z\| >5 | RMSZ | #\|Z\| >5 |
| 1 | A | 0.29 | 0/8850 | 0.53 | 0/11981 |
| 2 | B | 0.30 | 0/2652 | 0.55 | 1/3587 (0.0%) |
| All | All | 0.29 | 0/11502 | 0.54 | 1/15568 (0.0%) |

Chiral center outliers are detected by calculating the chiral volume of a chiral center and verifying if the center is modelled as a planar moiety or with the opposite hand.A planarity outlier is detected by checking planarity of atoms in a peptide group, atoms in a mainchain group or atoms of a sidechain that are expected to be planar.

| Mol | Chain | #Chirality outliers | #Planarity outliers |
|---|---|---|---|
| 1 | A | 0 | 1 |

There are no bond length outliers.

All (1) bond angle outliers are listed below:

| Mol | Chain | Res | Type | Atoms | Z | Observed($^o$) | Ideal($^o$) |
|---|---|---|---|---|---|---|---|
| 2 | B | 1168 | SER | N-CA-CB | 6.01 | 119.51 | 110.50 |

There are no chirality outliers.

All (1) planarity outliers are listed below:

| Mol | Chain | Res | Type | Group |
|---|---|---|---|---|
| 1 | A | 563 | ASP | Peptide |

#### 5.2 Too-close contacts (i)

In the following table, the Non-H and H(model) columns list the number of non-hydrogen atoms and hydrogen atoms in the chain respectively. The H(added) column lists the number of hydrogen atoms added and optimized by MolProbity. The Clashes column lists the number of clashes within the asymmetric unit, whereas Symm-Clashes lists symmetry related clashes.

CONFIDENTIAL VALIDATION REPORT

| Mol | Chain | Non-H | H(model) | H(added) | Clashes | Symm-Clashes |
|---|---|---|---|---|---|---|
| 1 | A | 8689 | 0 | 8671 | 148 | 1 |
| 2 | B | 2590 | 0 | 2507 | 43 | 0 |
| 3 | A | 9 | 0 | 0 | 0 | 0 |
| 3 | B | 1 | 0 | 0 | 0 | 0 |
| All | All | 11289 | 0 | 11178 | 188 | 1 |

The all-atom clashscore is defined as the number of clashes found per 1000 atoms (including hydrogen atoms). The all-atom clashscore for this structure is 8.

All (188) close contacts within the same asymmetric unit are listed below, sorted by their clash magnitude.

| Atom-1 | Atom-2 | Interatomic distance (Å) | Clash overlap (Å) |
|---|---|---|---|
| 1:A:969:GLU:HG2 | 1:A:971:ALA:H | 1.41 | 0.84 |
| 1:A:517:TYR:HE1 | 1:A:531:HIS:HD1 | 1.33 | 0.76 |
| 2:B:1120:TYR:CE1 | 2:B:1127:GLU:HG2 | 2.24 | 0.73 |
| 1:A:114:ARG:NH1 | 1:A:1079:GLU:OE1 | 2.21 | 0.73 |
| 2:B:1198:ASP:OD2 | 2:B:1222:ASN:ND2 | 2.21 | 0.73 |
| 1:A:1109:VAL:HG21 | 1:A:1126:ALA:HB2 | 1.71 | 0.72 |
| 1:A:36:ASN:ND2 | 1:A:1002:GLU:OE2 | 2.20 | 0.70 |
| 1:A:262:ASN:HD21 | 1:A:316:TYR:H | 1.39 | 0.69 |
| 1:A:262:ASN:ND2 | 1:A:316:TYR:H | 1.90 | 0.69 |
| 1:A:1044:SER:HG | 1:A:1047:TRP:HD1 | 1.40 | 0.69 |
| 2:B:1077:PHE:CD2 | 2:B:1307:VAL:HG21 | 2.27 | 0.68 |
| 1:A:467:GLN:HE22 | 1:A:524:GLN:H | 1.40 | 0.67 |
| 1:A:872:SER:HB3 | 1:A:914:LEU:HD13 | 1.79 | 0.63 |
| 1:A:307:GLU:OE2 | 1:A:383:LYS:NZ | 2.31 | 0.63 |
| 1:A:1055:GLN:HG3 | 1:A:1093:LEU:HD23 | 1.81 | 0.63 |
| 1:A:756:ALA:HB1 | 1:A:801:VAL:HG21 | 1.79 | 0.63 |
| 1:A:934:ALA:HB2 | 1:A:945:ILE:HD11 | 1.79 | 0.62 |
| 2:B:1259:ASN:OD1 | 2:B:1260:MET:N | 2.31 | 0.62 |
| 1:A:958:GLU:OE2 | 1:A:1009:HIS:NE2 | 2.33 | 0.62 |
| 1:A:263:ARG:HG2 | 1:A:271:TYR:CE1 | 2.35 | 0.62 |
| 1:A:842:GLU:OE1 | 2:B:1079:ARG:NH2 | 2.29 | 0.62 |
| 1:A:925:ASP:HB3 | 1:A:928:ARG:O | 2.01 | 0.61 |
| 1:A:213:GLU:HG2 | 1:A:215:GLU:H | 1.63 | 0.61 |
| 1:A:789:HIS:CD2 | 1:A:812:TYR:HA | 2.36 | 0.60 |
| 1:A:741:GLU:HG2 | 1:A:751:ALA:HA | 1.84 | 0.59 |
| 1:A:944:GLU:OE2 | 1:A:947:ARG:NH2 | 2.36 | 0.59 |
| 2:B:1120:TYR:HE1 | 2:B:1127:GLU:HG2 | 1.68 | 0.59 |
| 1:A:917:LYS:HG3 | 1:A:918:GLY:H | 1.67 | 0.58 |
| 1:A:517:TYR:HE1 | 1:A:531:HIS:ND1 | 2.02 | 0.58 |
| 1:A:59:GLY:HA2 | 1:A:1073:TRP:CZ3 | 2.38 | 0.58 |

*Continued on next page...*

CONFIDENTIAL VALIDATION REPORT

*Continued from previous page...*

| Atom-1 | Atom-2 | Interatomic distance (Å) | Clash overlap (Å) |
|---|---|---|---|
| 2:B:1077:PHE:CE2 | 2:B:1307:VAL:HG21 | 2.39 | 0.58 |
| 2:B:1301:PHE:CE1 | 2:B:1308:MET:HG2 | 2.39 | 0.58 |
| 1:A:301:ARG:NH1 | 1:A:303:GLU:OE2 | 2.24 | 0.57 |
| 2:B:1268:PRO:HD3 | 2:B:1301:PHE:CE2 | 2.39 | 0.57 |
| 1:A:72:GLU:OE2 | 1:A:103:ARG:NH2 | 2.36 | 0.57 |
| 1:A:1044:SER:OG | 1:A:1047:TRP:HD1 | 1.87 | 0.56 |
| 1:A:614:PHE:HE1 | 1:A:626:ARG:HG3 | 1.70 | 0.56 |
| 1:A:629:VAL:HG21 | 1:A:668:PHE:HE2 | 1.70 | 0.56 |
| 1:A:518:TYR:HD2 | 1:A:529:ILE:HB | 1.69 | 0.56 |
| 1:A:412:PRO:HD3 | 1:A:680:CYS:HB2 | 1.88 | 0.56 |
| 1:A:328:LEU:HB3 | 1:A:381:ALA:HB3 | 1.87 | 0.56 |
| 1:A:407:ILE:HG12 | 1:A:429:PHE:HE1 | 1.70 | 0.56 |
| 1:A:655:ARG:NH2 | 1:A:1138:ARG:HD3 | 2.21 | 0.56 |
| 1:A:340:SER:HB3 | 1:A:346:TYR:HE1 | 1.72 | 0.54 |
| 1:A:188:ARG:NH1 | 1:A:216:ALA:O | 2.37 | 0.54 |
| 1:A:642:ARG:HG2 | 1:A:647:THR:HG22 | 1.90 | 0.53 |
| 1:A:884:ILE:HG22 | 1:A:885:ASN:H | 1.74 | 0.53 |
| 2:B:1352:ARG:NH1 | 2:B:1385:ARG:HH22 | 2.07 | 0.53 |
| 1:A:81:THR:HG22 | 1:A:85:ASN:H | 1.73 | 0.53 |
| 1:A:969:GLU:OE2 | 1:A:973:ASN:HB2 | 2.09 | 0.53 |
| 2:B:1167:LYS:HA | 2:B:1170:PHE:HB2 | 1.90 | 0.53 |
| 1:A:563:ASP:OD2 | 1:A:567:ARG:NH2 | 2.27 | 0.53 |
| 2:B:1275:ILE:HG22 | 2:B:1276:ASN:H | 1.74 | 0.53 |
| 1:A:84:TYR:HE1 | 1:A:135:LEU:HD22 | 1.74 | 0.53 |
| 1:A:467:GLN:HE22 | 1:A:523:PRO:HA | 1.74 | 0.53 |
| 1:A:744:ASP:H | 1:A:749:THR:N | 2.06 | 0.52 |
| 1:A:38:ARG:NH2 | 1:A:54:GLU:OE1 | 2.41 | 0.52 |
| 1:A:629:VAL:HG21 | 1:A:668:PHE:CE2 | 2.44 | 0.52 |
| 1:A:564:ILE:HG23 | 1:A:588:PRO:HD3 | 1.92 | 0.52 |
| 1:A:417:PRO:HB3 | 1:A:481:GLN:HG2 | 1.90 | 0.52 |
| 1:A:738:SER:HB3 | 1:A:789:HIS:CE1 | 2.45 | 0.52 |
| 1:A:844:LYS:HG3 | 1:A:845:GLN:HG3 | 1.90 | 0.51 |
| 1:A:917:LYS:HG3 | 1:A:918:GLY:N | 2.26 | 0.51 |
| 1:A:285:LEU:HD22 | 1:A:297:LEU:HD21 | 1.93 | 0.51 |
| 2:B:1077:PHE:HD2 | 2:B:1307:VAL:HG21 | 1.73 | 0.51 |
| 2:B:1352:ARG:HH12 | 2:B:1385:ARG:HH22 | 1.58 | 0.51 |
| 1:A:252:ILE:HD12 | 1:A:252:ILE:H | 1.76 | 0.51 |
| 2:B:1136:SER:HB2 | 2:B:1154:ALA:HB1 | 1.93 | 0.51 |
| 2:B:1268:PRO:HD3 | 2:B:1301:PHE:HE2 | 1.76 | 0.51 |
| 1:A:889:ARG:HD2 | 1:A:891:TYR:CZ | 2.46 | 0.50 |
| 1:A:762:SER:O | 1:A:803:HIS:ND1 | 2.44 | 0.50 |

*Continued on next page...*

CONFIDENTIAL VALIDATION REPORT

*Continued from previous page...*

| Atom-1 | Atom-2 | Interatomic distance (Å) | Clash overlap (Å) |
|---|---|---|---|
| 2:B:1370:GLU:OE2 | 2:B:1385:ARG:NH1 | 2.45 | 0.50 |
| 1:A:397:HIS:NE2 | 1:A:705:ASP:OD1 | 2.40 | 0.50 |
| 1:A:836:VAL:HG13 | 2:B:1049:ASN:ND2 | 2.27 | 0.50 |
| 1:A:945:ILE:O | 1:A:990:GLN:HG2 | 2.11 | 0.50 |
| 2:B:1226:ASN:HA | 2:B:1263:SER:HB3 | 1.92 | 0.50 |
| 1:A:467:GLN:NE2 | 1:A:524:GLN:H | 2.10 | 0.50 |
| 1:A:743:GLN:HG3 | 1:A:782:PHE:CD2 | 2.47 | 0.49 |
| 2:B:1177:THR:HG23 | 2:B:1178:GLU:HG3 | 1.95 | 0.49 |
| 1:A:480:SER:HG | 1:A:485:ALA:H | 1.60 | 0.49 |
| 1:A:922:LEU:HD12 | 1:A:931:LEU:O | 2.13 | 0.49 |
| 1:A:44:VAL:HG21 | 1:A:317:LEU:HD22 | 1.93 | 0.49 |
| 2:B:1359:THR:HG22 | 2:B:1366:LEU:HD13 | 1.95 | 0.49 |
| 1:A:968:ALA:HB2 | 1:A:974:LEU:HD23 | 1.94 | 0.49 |
| 1:A:55:VAL:HG11 | 1:A:1065:VAL:HG21 | 1.94 | 0.49 |
| 1:A:1109:VAL:HG12 | 1:A:1129:LEU:HD12 | 1.94 | 0.49 |
| 1:A:235:GLU:HG3 | 1:A:254:LYS:HE2 | 1.94 | 0.48 |
| 1:A:263:ARG:HH12 | 1:A:266:PRO:HA | 1.78 | 0.48 |
| 2:B:1287:LEU:HD11 | 2:B:1290:THR:HG23 | 1.95 | 0.48 |
| 1:A:381:ALA:O | 1:A:385:GLY:N | 2.46 | 0.48 |
| 1:A:913:TYR:C | 1:A:914:LEU:HD12 | 2.34 | 0.48 |
| 1:A:742:VAL:O | 1:A:750:THR:N | 2.47 | 0.48 |
| 1:A:888:VAL:HB | 1:A:905:HIS:HB3 | 1.94 | 0.48 |
| 1:A:929:SER:OG | 1:A:948:ASP:HB3 | 2.13 | 0.48 |
| 1:A:402:ILE:HG22 | 1:A:404:LEU:HG | 1.96 | 0.48 |
| 1:A:894:THR:OG1 | 1:A:898:GLU:N | 2.41 | 0.48 |
| 1:A:4:ASN:HB2 | 1:A:1088:PHE:CD1 | 2.49 | 0.48 |
| 1:A:998:PHE:CZ | 1:A:1074:ARG:HD2 | 2.49 | 0.47 |
| 2:B:1337:PHE:CD1 | 2:B:1344:PRO:HA | 2.49 | 0.47 |
| 1:A:234:GLN:HG3 | 1:A:257:THR:HG22 | 1.96 | 0.47 |
| 1:A:268:GLY:O | 1:A:285:LEU:HD12 | 2.14 | 0.47 |
| 1:A:481:GLN:O | 1:A:484:LYS:NZ | 2.45 | 0.47 |
| 1:A:614:PHE:CE1 | 1:A:626:ARG:HG3 | 2.48 | 0.47 |
| 1:A:415:SER:O | 1:A:481:GLN:NE2 | 2.47 | 0.47 |
| 1:A:741:GLU:OE1 | 1:A:749:THR:HB | 2.15 | 0.47 |
| 2:B:1163:LEU:HB2 | 2:B:1176:PHE:HE2 | 1.79 | 0.47 |
| 1:A:490:TRP:CH2 | 1:A:517:TYR:HD2 | 2.33 | 0.47 |
| 2:B:1279:ILE:HD13 | 2:B:1289:HIS:HB2 | 1.96 | 0.47 |
| 1:A:617:ASN:O | 1:A:621:GLY:N | 2.38 | 0.46 |
| 2:B:1204:ASP:OD1 | 2:B:1205:ILE:N | 2.48 | 0.46 |
| 1:A:578:HIS:NE2 | 1:A:580:GLU:OE2 | 2.48 | 0.46 |
| 1:A:881:LEU:HD23 | 1:A:890:LEU:HD13 | 1.97 | 0.46 |

*Continued on next page...*

CONFIDENTIAL VALIDATION REPORT

*Continued from previous page...*

| Atom-1 | Atom-2 | Interatomic distance (Å) | Clash overlap (Å) |
|---|---|---|---|
| 1:A:518:TYR:CD2 | 1:A:529:ILE:HB | 2.50 | 0.46 |
| 1:A:286:GLU:O | 1:A:298:LYS:N | 2.45 | 0.46 |
| 1:A:396:ILE:HD13 | 1:A:693:LEU:HD12 | 1.97 | 0.46 |
| 1:A:315:THR:OG1 | 1:A:323:PHE:HB3 | 2.16 | 0.46 |
| 1:A:921:ILE:HB | 1:A:933:LEU:HB2 | 1.98 | 0.46 |
| 1:A:385:GLY:HA3 | 1:A:719:GLU:O | 2.16 | 0.46 |
| 1:A:1097:PHE:O | 1:A:1100:ILE:HG22 | 2.15 | 0.46 |
| 1:A:118:THR:OG1 | 1:A:134:ARG:NH2 | 2.48 | 0.46 |
| 2:B:1178:GLU:HB3 | 2:B:1196:LYS:HD2 | 1.97 | 0.46 |
| 2:B:1272:GLU:OE1 | 2:B:1342:TYR:OH | 2.26 | 0.46 |
| 1:A:1058:LEU:HD21 | 1:A:1097:PHE:HD1 | 1.80 | 0.46 |
| 1:A:413:LEU:HD21 | 1:A:468:LEU:HD21 | 1.97 | 0.46 |
| 1:A:84:TYR:CE1 | 1:A:135:LEU:HD22 | 2.51 | 0.46 |
| 1:A:282:MET:HB2 | 1:A:305:LEU:HD11 | 1.98 | 0.45 |
| 2:B:1297:CYS:HA | 2:B:1311:ALA:O | 2.17 | 0.45 |
| 1:A:637:VAL:HB | 1:A:652:CYS:HB2 | 1.98 | 0.45 |
| 1:A:492:GLU:HG3 | 1:A:512:VAL:HG21 | 1.99 | 0.44 |
| 1:A:467:GLN:NE2 | 1:A:523:PRO:HA | 2.31 | 0.44 |
| 1:A:947:ARG:HD2 | 1:A:949:PHE:CE1 | 2.52 | 0.44 |
| 1:A:917:LYS:HG2 | 1:A:962:ASP:OD1 | 2.18 | 0.44 |
| 2:B:1308:MET:HE3 | 2:B:1337:PHE:HB2 | 1.99 | 0.44 |
| 1:A:928:ARG:HH21 | 2:B:1364:CYS:HA | 1.83 | 0.44 |
| 1:A:182:TYR:OH | 1:A:209:GLN:OE1 | 2.31 | 0.44 |
| 1:A:560:LEU:HD13 | 1:A:567:ARG:HH11 | 1.82 | 0.44 |
| 1:A:271:TYR:HB2 | 1:A:283:LEU:HB3 | 1.99 | 0.44 |
| 1:A:922:LEU:HD22 | 1:A:965:PHE:CD1 | 2.52 | 0.44 |
| 2:B:1309:TYR:CE2 | 2:B:1359:THR:HG21 | 2.53 | 0.44 |
| 1:A:31:LEU:HD23 | 1:A:49:LEU:HD21 | 2.00 | 0.44 |
| 1:A:922:LEU:HD22 | 1:A:965:PHE:HD1 | 1.83 | 0.44 |
| 2:B:1083:ILE:HG13 | 2:B:1084:SER:N | 2.32 | 0.44 |
| 1:A:191:LYS:HE2 | 1:A:193:TYR:CE1 | 2.53 | 0.44 |
| 1:A:170:LEU:HD11 | 1:A:179:CYS:HB2 | 2.00 | 0.44 |
| 2:B:1142:GLU:OE1 | 2:B:1183:GLU:HA | 2.18 | 0.43 |
| 2:B:1221:ASN:HB3 | 2:B:1257:LYS:HD2 | 1.99 | 0.43 |
| 1:A:387:LEU:HB2 | 1:A:715:VAL:HB | 2.00 | 0.43 |
| 1:A:790:ASN:HD22 | 1:A:806:GLN:HA | 1.83 | 0.43 |
| 1:A:1124:ALA:HB1 | 1:A:1128:ASP:HB2 | 2.01 | 0.43 |
| 1:A:1051:LEU:HB2 | 1:A:1089:ILE:HD13 | 1.99 | 0.43 |
| 1:A:1106:GLN:HA | 1:A:1109:VAL:HG22 | 2.00 | 0.43 |
| 1:A:614:PHE:HD2 | 1:A:623:LEU:HD13 | 1.83 | 0.42 |
| 2:B:1081:ARG:HD3 | 2:B:1392:GLN:O | 2.19 | 0.42 |

*Continued on next page...*

CONFIDENTIAL VALIDATION REPORT

*Continued from previous page...*

| Atom-1 | Atom-2 | Interatomic distance (Å) | Clash overlap (Å) |
|---|---|---|---|
| 1:A:870:VAL:HA | 1:A:883:SER:O | 2.20 | 0.42 |
| 1:A:504:ASN:OD1 | 1:A:505:SER:N | 2.47 | 0.42 |
| 2:B:1356:ASP:OD1 | 2:B:1357:LEU:N | 2.49 | 0.42 |
| 1:A:392:ASN:HA | 1:A:710:LEU:HD23 | 2.00 | 0.42 |
| 1:A:659:ILE:HG23 | 1:A:668:PHE:HE1 | 1.84 | 0.42 |
| 2:B:1097:THR:N | 2:B:1111:GLY:O | 2.42 | 0.42 |
| 1:A:273:LEU:HB2 | 1:A:281:PHE:HB2 | 2.00 | 0.42 |
| 1:A:479:VAL:HG12 | 1:A:480:SER:O | 2.20 | 0.42 |
| 1:A:695:ASN:OD1 | 1:A:698:THR:N | 2.41 | 0.42 |
| 1:A:6:VAL:HG22 | 1:A:1040:VAL:HG22 | 2.02 | 0.42 |
| 1:A:90:GLU:HB3 | 1:A:101:ILE:CG2 | 2.49 | 0.42 |
| 2:B:1313:LEU:HD23 | 2:B:1314:GLN:N | 2.35 | 0.42 |
| 1:A:490:TRP:CH2 | 1:A:517:TYR:CD2 | 3.08 | 0.42 |
| 1:A:519:LEU:HD23 | 1:A:528:GLN:HA | 2.01 | 0.42 |
| 1:A:977:CYS:HB3 | 1:A:992:LEU:HB3 | 2.01 | 0.41 |
| 2:B:1226:ASN:ND2 | 2:B:1239:ASN:OD1 | 2.49 | 0.41 |
| 1:A:1007:PHE:CD1 | 1:A:1030:PHE:HB3 | 2.54 | 0.41 |
| 1:A:262:ASN:OD1 | 1:A:263:ARG:N | 2.53 | 0.41 |
| 1:A:879:LYS:NZ | 1:A:935:TYR:OH | 2.53 | 0.41 |
| 2:B:1258:PHE:HD2 | 2:B:1278:GLU:CD | 2.24 | 0.41 |
| 1:A:341:ASN:OD1 | 1:A:345:SER:N | 2.53 | 0.41 |
| 1:A:417:PRO:HD3 | 1:A:481:GLN:HE21 | 1.84 | 0.41 |
| 1:A:592:LEU:HG | 1:A:594:THR:HG23 | 2.02 | 0.41 |
| 1:A:639:ARG:HD3 | 1:A:679:MET:O | 2.21 | 0.41 |
| 1:A:884:ILE:O | 1:A:885:ASN:C | 2.59 | 0.41 |
| 1:A:536:HIS:ND1 | 1:A:562:THR:HB | 2.36 | 0.41 |
| 1:A:24:THR:HA | 1:A:91:TYR:CD2 | 2.56 | 0.41 |
| 1:A:680:CYS:SG | 1:A:692:ALA:HB3 | 2.61 | 0.40 |
| 1:A:68:ARG:HB2 | 1:A:75:ASP:OD1 | 2.21 | 0.40 |
| 1:A:830:ILE:HG22 | 1:A:873:MET:HE1 | 2.03 | 0.40 |
| 2:B:1261:ASN:OD1 | 2:B:1296:GLN:NE2 | 2.49 | 0.40 |
| 1:A:1030:PHE:CZ | 1:A:1038:GLY:HA3 | 2.56 | 0.40 |
| 1:A:743:GLN:OE1 | 1:A:749:THR:N | 2.54 | 0.40 |
| 2:B:1275:ILE:HG22 | 2:B:1276:ASN:N | 2.36 | 0.40 |
| 1:A:477:ARG:NH2 | 1:A:486:LEU:HD22 | 2.36 | 0.40 |

All (1) symmetry-related close contacts are listed below. The label for Atom-2 includes the symmetry operator and encoded unit-cell translations to be applied.

| Atom-1 | Atom-2 | Interatomic distance (Å) | Clash overlap (Å) |
|---|---|---|---|
| 1:A:346:TYR:OH | 1:A:420:GLU:OE2[2_555] | 2.18 | 0.02 |

CONFIDENTIAL VALIDATION REPORT

#### 5.3 Torsion angles (i)

##### 5.3.1 Protein backbone (i)

In the following table, the Percentiles column shows the percent Ramachandran outliers of the chain as a percentile score with respect to all X-ray entries followed by that with respect to entries of similar resolution.

The Analysed column shows the number of residues for which the backbone conformation was analysed, and the total number of residues.

| Mol | Chain | Analysed | Favoured | Allowed | Outliers | Percentiles | |
|---|---|---|---|---|---|---|---|
| 1 | A | 1093/1142 (96%) | 1031 (94%) | 62 (6%) | 0 | 100 | 100 |
| 2 | B | 313/373 (84%) | 298 (95%) | 15 (5%) | 0 | 100 | 100 |
| All | All | 1406/1515 (93%) | 1329 (94%) | 77 (6%) | 0 | 100 | 100 |

There are no Ramachandran outliers to report.

##### 5.3.2 Protein sidechains (i)

In the following table, the Percentiles column shows the percent sidechain outliers of the chain as a percentile score with respect to all X-ray entries followed by that with respect to entries of similar resolution.

The Analysed column shows the number of residues for which the sidechain conformation was analysed, and the total number of residues.

| Mol | Chain | Analysed | Rotameric | Outliers | Percentiles | |
|---|---|---|---|---|---|---|
| 1 | A | 973/1000 (97%) | 972 (100%) | 1 (0%) | 93 | 98 |
| 2 | B | 285/327 (87%) | 285 (100%) | 0 | 100 | 100 |
| All | All | 1258/1327 (95%) | 1257 (100%) | 1 (0%) | 93 | 98 |

All (1) residues with a non-rotameric sidechain are listed below:

| Mol | Chain | Res | Type |
|---|---|---|---|
| 1 | A | 617 | ASN |

Some sidechains can be flipped to improve hydrogen bonding and reduce clashes. All (3) such sidechains are listed below:

| Mol | Chain | Res | Type |
|---|---|---|---|
| 1 | A | 467 | GLN |
| 1 | A | 481 | GLN |

*Continued on next page...*

CONFIDENTIAL VALIDATION REPORT

*Continued from previous page...*

| Mol | Chain | Res | Type |
|---|---|---|---|
| 1 | A | 617 | ASN |

##### 5.3.3 RNA (i)

There are no RNA molecules in this entry.

#### 5.4 Non-standard residues in protein, DNA, RNA chains (i)

There are no non-standard protein/DNA/RNA residues in this entry.

#### 5.5 Carbohydrates (i)

There are no monosaccharides in this entry.

#### 5.6 Ligand geometry (i)

There are no ligands in this entry.

#### 5.7 Other polymers (i)

There are no such residues in this entry.

#### 5.8 Polymer linkage issues (i)

There are no chain breaks in this entry.

CONFIDENTIAL VALIDATION REPORT

### 6 Fit of model and data (i)

#### 6.1 Protein, DNA and RNA chains (i)

In the following table, the column labelled '#RSRZ> 2' contains the number (and percentage) of RSRZ outliers, followed by percent RSRZ outliers for the chain as percentile scores relative to all X-ray entries and entries of similar resolution. The OWAB column contains the minimum, median, $95^{th}$ percentile and maximum values of the occupancy-weighted average B-factor per residue. The column labelled 'Q< 0.9' lists the number of (and percentage) of residues with an average occupancy less than 0.9.

| Mol | Chain | Analysed | <RSRZ> | #RSRZ>2 | | | OWAB(Å$^2$) | Q<0.9 |
|---|---|---|---|---|---|---|---|---|
| 1 | A | 1106/1142 (96%) | -0.09 | 7 (0%) | 89 | 78 | 69, 108, 160, 252 | 0 |
| 2 | B | 322/373 (86%) | -0.05 | 4 (1%) | 79 | 61 | 76, 114, 159, 189 | 0 |
| All | All | 1428/1515 (94%) | -0.08 | 11 (0%) | 86 | 72 | 69, 109, 160, 252 | 0 |

All (11) RSRZ outliers are listed below:

| Mol | Chain | Res | Type | RSRZ |
|---|---|---|---|---|
| 1 | A | 761 | LEU | 2.9 |
| 1 | A | 855 | ASP | 2.7 |
| 1 | A | 938 | MET | 2.5 |
| 2 | B | 1355 | PHE | 2.4 |
| 2 | B | 1176 | PHE | 2.3 |
| 2 | B | 1212 | LEU | 2.3 |
| 1 | A | 1129 | LEU | 2.2 |
| 1 | A | 297 | LEU | 2.2 |
| 1 | A | 1126 | ALA | 2.1 |
| 2 | B | 1214 | LEU | 2.1 |
| 1 | A | 1025 | GLN | 2.0 |

#### 6.2 Non-standard residues in protein, DNA, RNA chains (i)

There are no non-standard protein/DNA/RNA residues in this entry.

#### 6.3 Carbohydrates (i)

There are no monosaccharides in this entry.

CONFIDENTIAL VALIDATION REPORT

#### 6.4 Ligands (i)

There are no ligands in this entry.

#### 6.5 Other polymers (i)

There are no such residues in this entry.

CONFIDENTIAL VALIDATION REPORT

W O R L D W I D E PDB PROTEIN DATA BANK

### Full wwPDB X-ray Structure Validation Report ⓘ

Jul 31, 2020 – 03:52 PM BST

| | | |
|---|---|---|
| PDB ID | : | 6ZX9 |
| Title | : | Crystal structure of SIV Vpr,fused to T4 lysozyme, isolated from moustached monkey, bound to human DDB1 and human DCAF1 (amino acid residues 1046-1396) |
| Deposited on | : | 2020-07-29 |
| Resolution | : | 2.52 Å(reported) |

This is a Full wwPDB X-ray Structure Validation Report.

This report is produced by the wwPDB biocuration pipeline after annotation of the structure.

We welcome your comments at **
A user guide is available at
https://www.wwpdb.org/validation/2017/XrayValidationReportHelp
with specific help available everywhere you see the ⓘ symbol.

CONFIDENTIAL VALIDATION REPORT

---

The following versions of software and data (see references ⓘ) were used in the production of this report:

| | | |
|---|---|---|
| MolProbity | : | 4.02b-467 |
| Mogul | : | 1.8.5 (274361), CSD as541be (2020) |
| Xtriage (Phenix) | : | 1.13 |
| EDS | : | 2.13 |
| buster-report | : | 1.1.7 (2018) |
| Percentile statistics | : | 20191225.v01 (using entries in the PDB archive December 25th 2019) |
| Refmac | : | 5.8.0158 |
| CCP4 | : | 7.0.044 (Gargrove) |
| Ideal geometry (proteins) | : | Engh & Huber (2001) |
| Ideal geometry (DNA, RNA) | : | Parkinson et al. (1996) |
| Validation Pipeline (wwPDB-VP) | : | 2.13 |

### 1 Overall quality at a glance (i)

The following experimental techniques were used to determine the structure:
*X-RAY DIFFRACTION*

The reported resolution of this entry is 2.52 Å.

Percentile scores (ranging between 0-100) for global validation metrics of the entry are shown in the following graphic. The table shows the number of entries on which the scores are based.

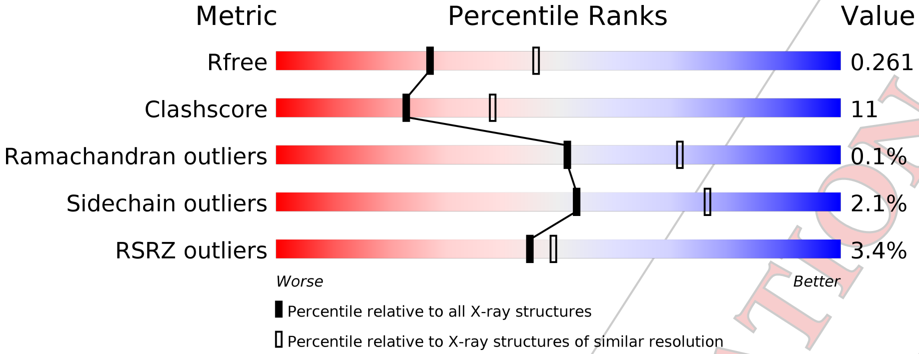

| Metric | Whole archive (#Entries) | Similar resolution (#Entries, resolution range(Å)) |
|---|---|---|
| $R_{free}$ | 130704 | 5743 (2.54-2.50) |
| Clashscore | 141614 | 6463 (2.54-2.50) |
| Ramachandran outliers | 138981 | 6335 (2.54-2.50) |
| Sidechain outliers | 138945 | 6337 (2.54-2.50) |
| RSRZ outliers | 127900 | 5630 (2.54-2.50) |

The table below summarises the geometric issues observed across the polymeric chains and their fit to the electron density. The red, orange, yellow and green segments on the lower bar indicate the fraction of residues that contain outliers for >=3, 2, 1 and 0 types of geometric quality criteria respectively. A grey segment represents the fraction of residues that are not modelled. The numeric value for each fraction is indicated below the corresponding segment, with a dot representing fractions <=5% The upper red bar (where present) indicates the fraction of residues that have poor fit to the electron density. The numeric value is given above the bar.

| Mol | Chain | Length | Quality of chain |
|---|---|---|---|
| 1 | A | 1142 | 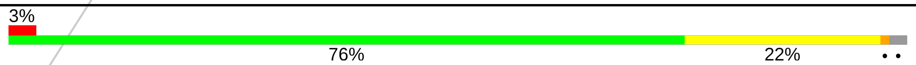  |
| 2 | B | 360 | 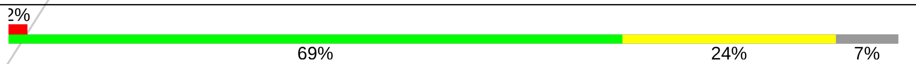  |
| 3 | C | 258 | 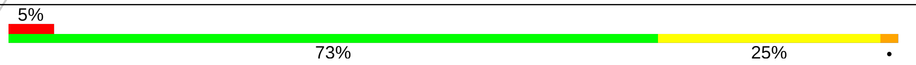  |

CONFIDENTIAL VALIDATION REPORT

#### 2 Entry composition (i)

There are 6 unique types of molecules in this entry. The entry contains 13858 atoms, of which 0 are hydrogens and 0 are deuteriums.

In the tables below, the ZeroOcc column contains the number of atoms modelled with zero occupancy, the AltConf column contains the number of residues with at least one atom in alternate conformation and the Trace column contains the number of residues modelled with at most 2 atoms.

- Molecule 1 is a protein called DNA damage-binding protein 1.

| Mol | Chain | Residues | Atoms | | | | | ZeroOcc | AltConf | Trace |
|---|---|---|---|---|---|---|---|---|---|---|
| | | | Total | C | N | O | S | | | |
| 1 | A | 1124 | 8808 | 5581 | 1485 | 1695 | 47 | 0 | 1 | 0 |

There are 2 discrepancies between the modelled and reference sequences:

| Chain | Residue | Modelled | Actual | Comment | Reference |
|---|---|---|---|---|---|
| A | -1 | GLY | - | expression tag | UNP Q16531 |
| A | 0 | SER | - | expression tag | UNP Q16531 |

- Molecule 2 is a protein called DDB1- and CUL4-associated factor 1.

| Mol | Chain | Residues | Atoms | | | | | ZeroOcc | AltConf | Trace |
|---|---|---|---|---|---|---|---|---|---|---|
| | | | Total | C | N | O | S | | | |
| 2 | B | 335 | 2670 | 1684 | 467 | 501 | 18 | 0 | 0 | 0 |

There are 9 discrepancies between the modelled and reference sequences:

| Chain | Residue | Modelled | Actual | Comment | Reference |
|---|---|---|---|---|---|
| B | 1045 | MET | - | initiating methionine | UNP Q9Y4B6 |
| B | 1397 | GLU | - | expression tag | UNP Q9Y4B6 |
| B | 1398 | LEU | - | expression tag | UNP Q9Y4B6 |
| B | 1399 | ALA | - | expression tag | UNP Q9Y4B6 |
| B | 1400 | LEU | - | expression tag | UNP Q9Y4B6 |
| B | 1401 | VAL | - | expression tag | UNP Q9Y4B6 |
| B | 1402 | PRO | - | expression tag | UNP Q9Y4B6 |
| B | 1403 | ARG | - | expression tag | UNP Q9Y4B6 |
| B | 1404 | GLY | - | expression tag | UNP Q9Y4B6 |

- Molecule 3 is a protein called Endolysin.

CONFIDENTIAL VALIDATION REPORT

| Mol | Chain | Residues | Atoms | | | | | ZeroOcc | AltConf | Trace |
|---|---|---|---|---|---|---|---|---|---|---|
| | | | Total | C | N | O | S | | | |
| 3 | C | 257 | 2078 | 1306 | 384 | 379 | 9 | 0 | 0 | 0 |

There are 98 discrepancies between the modelled and reference sequences:

| Chain | Residue | Modelled | Actual | Comment | Reference |
|---|---|---|---|---|---|
| C | -156 | HIS | GLU | conflict | UNP A0A097J809 |
| C | -113 | THR | CYS | conflict | UNP A0A097J809 |
| C | -70 | ALA | CYS | conflict | UNP A0A097J809 |
| C | -2 | ALA | - | expression tag | UNP A0A097J809 |
| C | -1 | ALA | - | expression tag | UNP A0A097J809 |
| C | 0 | ALA | - | expression tag | UNP A0A097J809 |
| C | 1 | MET | - | expression tag | UNP A0A097J809 |
| C | 2 | GLU | - | expression tag | UNP A0A097J809 |
| C | 3 | ARG | - | expression tag | UNP A0A097J809 |
| C | 4 | VAL | - | expression tag | UNP A0A097J809 |
| C | 5 | PRO | - | expression tag | UNP A0A097J809 |
| C | 6 | PRO | - | expression tag | UNP A0A097J809 |
| C | 7 | SER | - | expression tag | UNP A0A097J809 |
| C | 8 | HIS | - | expression tag | UNP A0A097J809 |
| C | 9 | ARG | - | expression tag | UNP A0A097J809 |
| C | 10 | PRO | - | expression tag | UNP A0A097J809 |
| C | 11 | PRO | - | expression tag | UNP A0A097J809 |
| C | 12 | TRP | - | expression tag | UNP A0A097J809 |
| C | 13 | HIS | - | expression tag | UNP A0A097J809 |
| C | 14 | SER | - | expression tag | UNP A0A097J809 |
| C | 15 | ARG | - | expression tag | UNP A0A097J809 |
| C | 16 | VAL | - | expression tag | UNP A0A097J809 |
| C | 17 | VAL | - | expression tag | UNP A0A097J809 |
| C | 18 | PRO | - | expression tag | UNP A0A097J809 |
| C | 19 | THR | - | expression tag | UNP A0A097J809 |
| C | 20 | THR | - | expression tag | UNP A0A097J809 |
| C | 21 | MET | - | expression tag | UNP A0A097J809 |
| C | 22 | GLN | - | expression tag | UNP A0A097J809 |
| C | 23 | GLN | - | expression tag | UNP A0A097J809 |
| C | 24 | ALA | - | expression tag | UNP A0A097J809 |
| C | 25 | GLN | - | expression tag | UNP A0A097J809 |
| C | 26 | GLN | - | expression tag | UNP A0A097J809 |
| C | 27 | ALA | - | expression tag | UNP A0A097J809 |
| C | 28 | MET | - | expression tag | UNP A0A097J809 |
| C | 29 | TRP | - | expression tag | UNP A0A097J809 |
| C | 30 | ASP | - | expression tag | UNP A0A097J809 |
| C | 31 | LEU | - | expression tag | UNP A0A097J809 |

*Continued on next page...*

CONFIDENTIAL VALIDATION REPORT

*Continued from previous page...*

| Chain | Residue | Modelled | Actual | Comment | Reference |
|---|---|---|---|---|---|
| C | 32 | ASN | - | expression tag | UNP A0A097J809 |
| C | 33 | GLU | - | expression tag | UNP A0A097J809 |
| C | 34 | GLU | - | expression tag | UNP A0A097J809 |
| C | 35 | ALA | - | expression tag | UNP A0A097J809 |
| C | 36 | GLU | - | expression tag | UNP A0A097J809 |
| C | 37 | LYS | - | expression tag | UNP A0A097J809 |
| C | 38 | HIS | - | expression tag | UNP A0A097J809 |
| C | 39 | PHE | - | expression tag | UNP A0A097J809 |
| C | 40 | SER | - | expression tag | UNP A0A097J809 |
| C | 41 | ARG | - | expression tag | UNP A0A097J809 |
| C | 42 | GLU | - | expression tag | UNP A0A097J809 |
| C | 43 | GLU | - | expression tag | UNP A0A097J809 |
| C | 44 | LEU | - | expression tag | UNP A0A097J809 |
| C | 45 | ARG | - | expression tag | UNP A0A097J809 |
| C | 46 | GLY | - | expression tag | UNP A0A097J809 |
| C | 47 | ILE | - | expression tag | UNP A0A097J809 |
| C | 48 | TRP | - | expression tag | UNP A0A097J809 |
| C | 49 | ASN | - | expression tag | UNP A0A097J809 |
| C | 50 | ASP | - | expression tag | UNP A0A097J809 |
| C | 51 | VAL | - | expression tag | UNP A0A097J809 |
| C | 52 | THR | - | expression tag | UNP A0A097J809 |
| C | 53 | GLU | - | expression tag | UNP A0A097J809 |
| C | 54 | LEU | - | expression tag | UNP A0A097J809 |
| C | 55 | PRO | - | expression tag | UNP A0A097J809 |
| C | 56 | ALA | - | expression tag | UNP A0A097J809 |
| C | 57 | ASP | - | expression tag | UNP A0A097J809 |
| C | 58 | PRO | - | expression tag | UNP A0A097J809 |
| C | 59 | ASN | - | expression tag | UNP A0A097J809 |
| C | 60 | TRP | - | expression tag | UNP A0A097J809 |
| C | 61 | THR | - | expression tag | UNP A0A097J809 |
| C | 62 | VAL | - | expression tag | UNP A0A097J809 |
| C | 63 | ASP | - | expression tag | UNP A0A097J809 |
| C | 64 | GLN | - | expression tag | UNP A0A097J809 |
| C | 65 | ALA | - | expression tag | UNP A0A097J809 |
| C | 66 | ALA | - | expression tag | UNP A0A097J809 |
| C | 67 | ILE | - | expression tag | UNP A0A097J809 |
| C | 68 | ALA | - | expression tag | UNP A0A097J809 |
| C | 69 | CYS | - | expression tag | UNP A0A097J809 |
| C | 70 | ALA | - | expression tag | UNP A0A097J809 |
| C | 71 | ILE | - | expression tag | UNP A0A097J809 |
| C | 72 | ASP | - | expression tag | UNP A0A097J809 |
| C | 73 | TYR | - | expression tag | UNP A0A097J809 |

*Continued on next page...*

CONFIDENTIAL VALIDATION REPORT

*Continued from previous page...*

| Chain | Residue | Modelled | Actual | Comment | Reference |
|---|---|---|---|---|---|
| C | 74 | ILE | - | expression tag | UNP A0A097J809 |
| C | 75 | ARG | - | expression tag | UNP A0A097J809 |
| C | 76 | ARG | - | expression tag | UNP A0A097J809 |
| C | 77 | THR | - | expression tag | UNP A0A097J809 |
| C | 78 | GLN | - | expression tag | UNP A0A097J809 |
| C | 79 | THR | - | expression tag | UNP A0A097J809 |
| C | 80 | LEU | - | expression tag | UNP A0A097J809 |
| C | 81 | LEU | - | expression tag | UNP A0A097J809 |
| C | 82 | PHE | - | expression tag | UNP A0A097J809 |
| C | 83 | ARG | - | expression tag | UNP A0A097J809 |
| C | 84 | HIS | - | expression tag | UNP A0A097J809 |
| C | 85 | TYR | - | expression tag | UNP A0A097J809 |
| C | 86 | ARG | - | expression tag | UNP A0A097J809 |
| C | 87 | GLU | - | expression tag | UNP A0A097J809 |
| C | 88 | GLY | - | expression tag | UNP A0A097J809 |
| C | 89 | CYS | - | expression tag | UNP A0A097J809 |
| C | 90 | TYR | - | expression tag | UNP A0A097J809 |
| C | 91 | HIS | - | expression tag | UNP A0A097J809 |
| C | 92 | ARG | - | expression tag | UNP A0A097J809 |

- Molecule 4 is GLYCEROL (three-letter code: GOL) (formula: $C_3H_8O_3$).

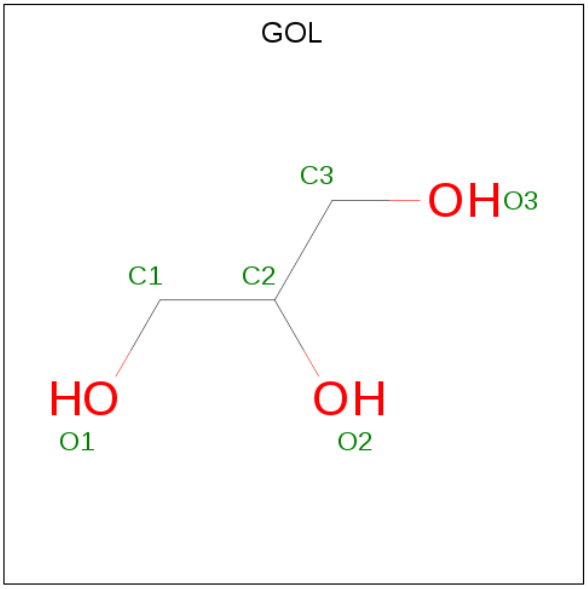

| Mol | Chain | Residues | Atoms | ZeroOcc | AltConf |
|---|---|---|---|---|---|
| 4 | A | 1 | Total C O<br>6 3 3 | 0 | 0 |
| 4 | A | 1 | Total C O<br>6 3 3 | 0 | 0 |

*Continued on next page...*

CONFIDENTIAL VALIDATION REPORT

*Continued from previous page...*

| Mol | Chain | Residues | Atoms | ZeroOcc | AltConf |
|---|---|---|---|---|---|
| 4 | A | 1 | Total C O<br>6 3 3 | 0 | 0 |
| 4 | A | 1 | Total C O<br>6 3 3 | 0 | 0 |
| 4 | A | 1 | Total C O<br>6 3 3 | 0 | 0 |
| 4 | A | 1 | Total C O<br>6 3 3 | 0 | 0 |
| 4 | B | 1 | Total C O<br>6 3 3 | 0 | 0 |
| 4 | C | 1 | Total C O<br>6 3 3 | 0 | 0 |

- Molecule 5 is ZINC ION (three-letter code: ZN) (formula: Zn) (labeled as "Ligand of Interest" by author).

| Mol | Chain | Residues | Atoms | ZeroOcc | AltConf |
|---|---|---|---|---|---|
| 5 | C | 1 | Total Zn<br>1 1 | 0 | 0 |

- Molecule 6 is water.

| Mol | Chain | Residues | Atoms | ZeroOcc | AltConf |
|---|---|---|---|---|---|
| 6 | A | 202 | Total O<br>202 202 | 0 | 0 |
| 6 | B | 37 | Total O<br>37 37 | 0 | 0 |
| 6 | C | 14 | Total O<br>14 14 | 0 | 0 |

CONFIDENTIAL VALIDATION REPORT

### 3 Residue-property plots

These plots are drawn for all protein, RNA, DNA and oligosaccharide chains in the entry. The first graphic for a chain summarises the proportions of the various outlier classes displayed in the second graphic. The second graphic shows the sequence view annotated by issues in geometry and electron density. Residues are color-coded according to the number of geometric quality criteria for which they contain at least one outlier: green = 0, yellow = 1, orange = 2 and red = 3 or more. A red dot above a residue indicates a poor fit to the electron density (RSRZ > 2). Stretches of 2 or more consecutive residues without any outlier are shown as a green connector. Residues present in the sample, but not in the model, are shown in grey.

- Molecule 1: DNA damage-binding protein 1

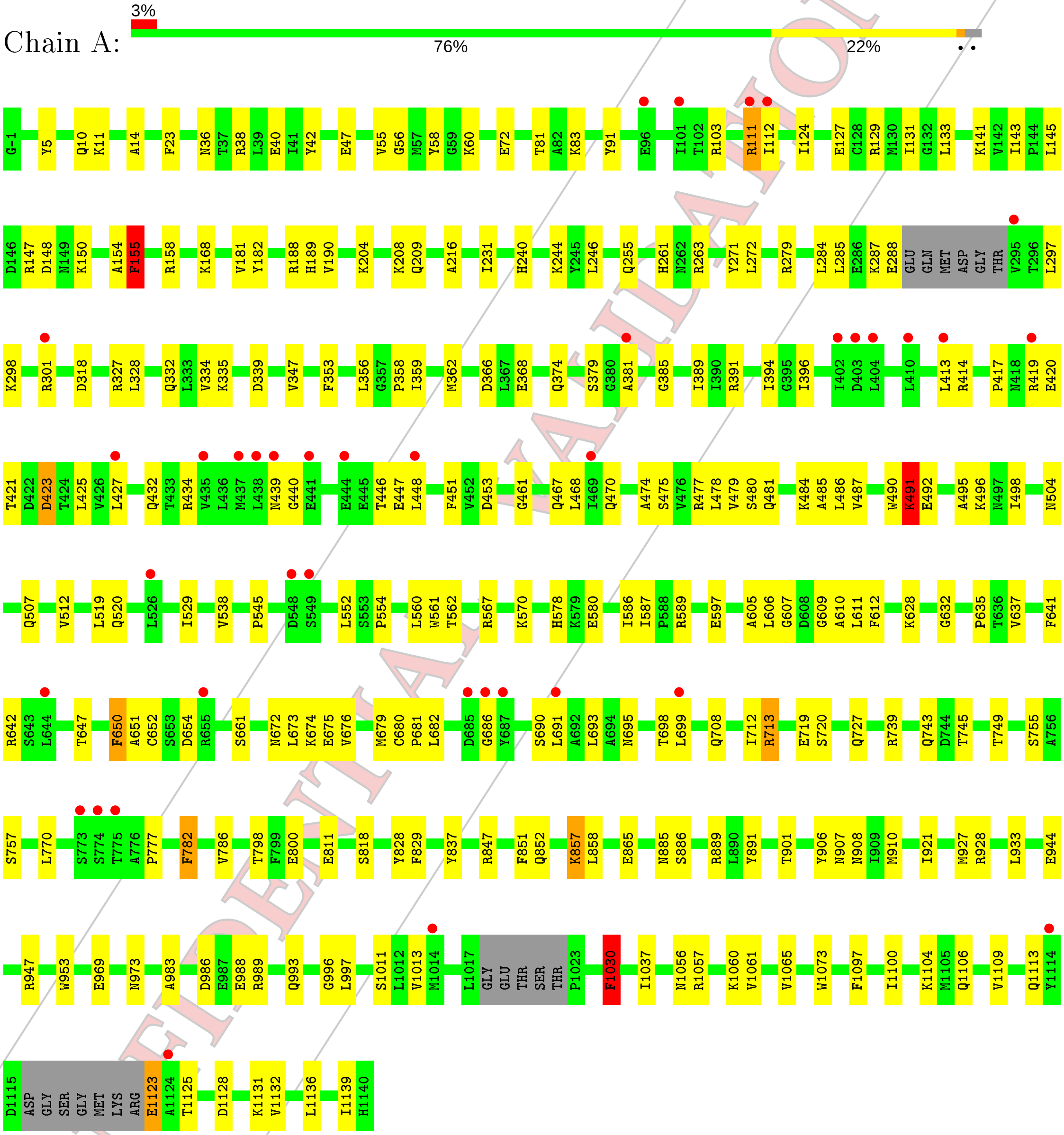

- Molecule 2: DDB1- and CUL4-associated factor 1

CONFIDENTIAL VALIDATION REPORT

Chain B:

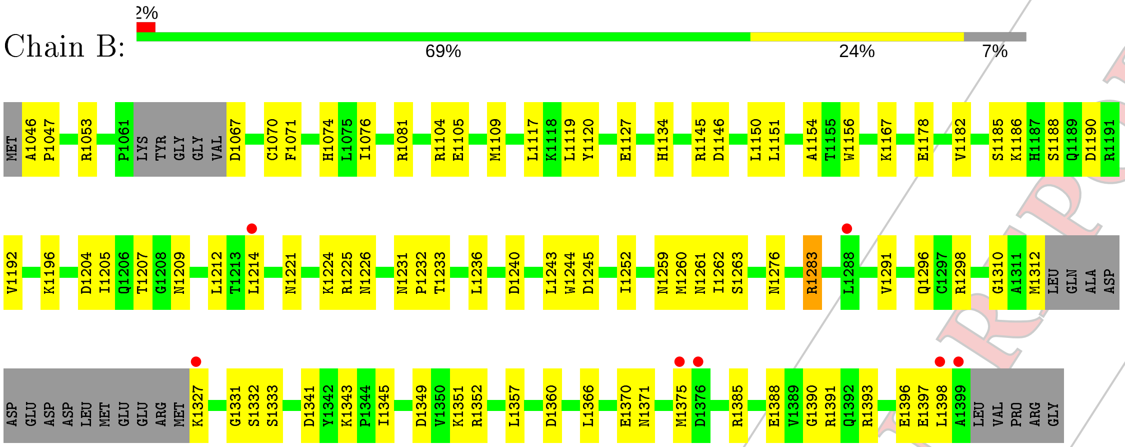

• Molecule 3: Endolysin

Chain C:

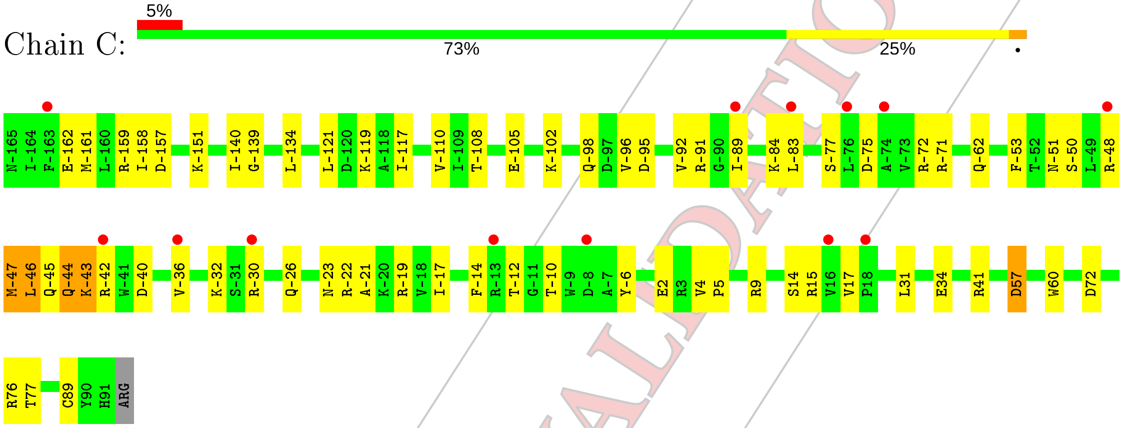

CONFIDENTIAL VALIDATION REPORT

### 4 Data and refinement statistics (i)

| Property | Value | Source |
|---|---|---|
| Space group | P 21 21 2 | Depositor |
| Cell constants<br>a, b, c, $\alpha$, $\beta$, $\gamma$ | 265.90Å 95.54Å 98.35Å<br>90.00° 90.00° 90.00° | Depositor |
| Resolution (Å) | 79.07 – 2.52<br>79.07 – 2.52 | Depositor<br>EDS |
| % Data completeness<br>(in resolution range) | 99.2 (79.07-2.52)<br>89.0 (79.07-2.52) | Depositor<br>EDS |
| $R_{merge}$ | (Not available) | Depositor |
| $R_{sym}$ | (Not available) | Depositor |
| $< I/\sigma(I) >$ [1] | 0.81 (at 2.51Å) | Xtriage |
| Refinement program | PHENIX 1.12_2829, PHENIX 1.12_2829 | Depositor |
| R, $R_{free}$ | 0.216 , 0.260<br>0.216 , 0.261 | Depositor<br>DCC |
| $R_{free}$ test set | 3948 reflections (4.62%) | wwPDB-VP |
| Wilson B-factor (Å$^2$) | 46.3 | Xtriage |
| Anisotropy | 0.133 | Xtriage |
| Bulk solvent $k_{sol}$(e/Å$^3$), $B_{sol}$(Å$^2$) | 0.31 , 47.9 | EDS |
| L-test for twinning[2] | $< \|L\| > = 0.48$, $< L^2 > = 0.32$ | Xtriage |
| Estimated twinning fraction | 0.018 for -h,l,k | Xtriage |
| $F_o$,$F_c$ correlation | 0.93 | EDS |
| Total number of atoms | 13858 | wwPDB-VP |
| Average B, all atoms (Å$^2$) | 71.0 | wwPDB-VP |

Xtriage's analysis on translational NCS is as follows: *The largest off-origin peak in the Patterson function is 2.63% of the height of the origin peak. No significant pseudotranslation is detected.*

[1] Intensities estimated from amplitudes.

[2] Theoretical values of $< |L| >$, $< L^2 >$ for acentric reflections are 0.5, 0.333 respectively for untwinned datasets, and 0.375, 0.2 for perfectly twinned datasets.

CONFIDENTIAL VALIDATION REPORT

### 5 Model quality (i)

#### 5.1 Standard geometry (i)

Bond lengths and bond angles in the following residue types are not validated in this section: GOL, ZN

The Z score for a bond length (or angle) is the number of standard deviations the observed value is removed from the expected value. A bond length (or angle) with $|Z| > 5$ is considered an outlier worth inspection. RMSZ is the root-mean-square of all Z scores of the bond lengths (or angles).

| Mol | Chain | Bond lengths | | Bond angles | |
|---|---|---|---|---|---|
| | | RMSZ | #\|Z\| >5 | RMSZ | #\|Z\| >5 |
| 1 | A | 0.29 | 0/8973 | 0.58 | 3/12154 (0.0%) |
| 2 | B | 0.28 | 0/2731 | 0.50 | 0/3696 |
| 3 | C | 0.31 | 0/2124 | 0.48 | 0/2878 |
| All | All | 0.29 | 0/13828 | 0.55 | 3/18728 (0.0%) |

Chiral center outliers are detected by calculating the chiral volume of a chiral center and verifying if the center is modelled as a planar moiety or with the opposite hand.A planarity outlier is detected by checking planarity of atoms in a peptide group, atoms in a mainchain group or atoms of a sidechain that are expected to be planar.

| Mol | Chain | #Chirality outliers | #Planarity outliers |
|---|---|---|---|
| 1 | A | 0 | 1 |

There are no bond length outliers.

All (3) bond angle outliers are listed below:

| Mol | Chain | Res | Type | Atoms | Z | Observed($^o$) | Ideal($^o$) |
|---|---|---|---|---|---|---|---|
| 1 | A | 1030 | PHE | CB-CG-CD1 | 5.67 | 124.77 | 120.80 |
| 1 | A | 491 | LYS | CD-CE-NZ | 5.34 | 123.97 | 111.70 |
| 1 | A | 155 | PHE | CB-CG-CD1 | 5.03 | 124.32 | 120.80 |

There are no chirality outliers.

All (1) planarity outliers are listed below:

| Mol | Chain | Res | Type | Group |
|---|---|---|---|---|
| 1 | A | 928 | ARG | Peptide |

CONFIDENTIAL VALIDATION REPORT

#### 5.2 Too-close contacts (i)

In the following table, the Non-H and H(model) columns list the number of non-hydrogen atoms and hydrogen atoms in the chain respectively. The H(added) column lists the number of hydrogen atoms added and optimized by MolProbity. The Clashes column lists the number of clashes within the asymmetric unit, whereas Symm-Clashes lists symmetry related clashes.

| Mol | Chain | Non-H | H(model) | H(added) | Clashes | Symm-Clashes |
|---|---|---|---|---|---|---|
| 1 | A | 8808 | 0 | 8782 | 195 | 0 |
| 2 | B | 2670 | 0 | 2568 | 71 | 0 |
| 3 | C | 2078 | 0 | 2059 | 51 | 0 |
| 4 | A | 36 | 0 | 48 | 2 | 0 |
| 4 | B | 6 | 0 | 8 | 0 | 0 |
| 4 | C | 6 | 0 | 8 | 0 | 0 |
| 5 | C | 1 | 0 | 0 | 0 | 0 |
| 6 | A | 202 | 0 | 0 | 8 | 0 |
| 6 | B | 37 | 0 | 0 | 1 | 0 |
| 6 | C | 14 | 0 | 0 | 2 | 0 |
| All | All | 13858 | 0 | 13473 | 309 | 0 |

The all-atom clashscore is defined as the number of clashes found per 1000 atoms (including hydrogen atoms). The all-atom clashscore for this structure is 11.

All (309) close contacts within the same asymmetric unit are listed below, sorted by their clash magnitude.

| Atom-1 | Atom-2 | Interatomic distance (Å) | Clash overlap (Å) |
|---|---|---|---|
| 2:B:1393:ARG:HD2 | 2:B:1398:LEU:CD2 | 1.28 | 1.56 |
| 2:B:1393:ARG:CD | 2:B:1398:LEU:HD21 | 1.30 | 1.55 |
| 1:A:111:ARG:NH1 | 1:A:112:ILE:CG2 | 1.68 | 1.54 |
| 1:A:111:ARG:NH1 | 1:A:112:ILE:HG22 | 0.83 | 1.15 |
| 1:A:906:TYR:OH | 2:B:1396:GLU:OE1 | 1.71 | 1.07 |
| 2:B:1393:ARG:NE | 2:B:1398:LEU:HD21 | 1.77 | 0.98 |
| 1:A:1113:GLN:OE1 | 1:A:1123:GLU:HA | 1.66 | 0.94 |
| 2:B:1393:ARG:HD2 | 2:B:1398:LEU:HD23 | 1.55 | 0.88 |
| 1:A:491:LYS:HA | 1:A:491:LYS:HE2 | 1.56 | 0.85 |
| 2:B:1145:ARG:NH2 | 2:B:1186:LYS:O | 2.10 | 0.84 |
| 2:B:1393:ARG:HD2 | 2:B:1398:LEU:CG | 2.08 | 0.83 |
| 1:A:111:ARG:NH1 | 1:A:112:ILE:HG21 | 1.95 | 0.82 |
| 1:A:496:LYS:O | 6:A:1301:HOH:O | 1.96 | 0.82 |
| 2:B:1393:ARG:CD | 2:B:1398:LEU:CD2 | 2.14 | 0.81 |
| 2:B:1207:THR:HG23 | 2:B:1209:ASN:H | 1.46 | 0.81 |
| 1:A:467:GLN:HG3 | 1:A:478:LEU:HD11 | 1.61 | 0.79 |
| 3:C:-72:ARG:NH1 | 3:C:-14:PHE:O | 2.15 | 0.79 |

Continued on next page...

CONFIDENTIAL VALIDATION REPORT

*Continued from previous page...*

| Atom-1 | Atom-2 | Interatomic distance (Å) | Clash overlap (Å) |
|---|---|---|---|
| 1:A:111:ARG:HH11 | 1:A:112:ILE:HG22 | 1.00 | 0.79 |
| 3:C:-53:PHE:HB3 | 3:C:-50:SER:HB2 | 1.65 | 0.79 |
| 1:A:654:ASP:O | 1:A:675:GLU:HG2 | 1.82 | 0.77 |
| 3:C:-75:ASP:OD1 | 3:C:-72:ARG:HD2 | 1.86 | 0.75 |
| 3:C:-157:ASP:OD1 | 3:C:-19:ARG:NH1 | 2.20 | 0.74 |
| 1:A:606:LEU:HD11 | 1:A:612:PHE:HE1 | 1.53 | 0.74 |
| 1:A:11:LYS:NZ | 1:A:38:ARG:HH21 | 1.87 | 0.73 |
| 1:A:285:LEU:HB3 | 1:A:297:LEU:HD11 | 1.71 | 0.72 |
| 2:B:1259:ASN:HD21 | 2:B:1296:GLN:HE21 | 1.37 | 0.72 |
| 3:C:14:SER:HB3 | 3:C:17:VAL:HG12 | 1.72 | 0.72 |
| 2:B:1370:GLU:OE1 | 2:B:1385:ARG:NH1 | 2.23 | 0.71 |
| 1:A:244:LYS:HE2 | 1:A:246:LEU:HD11 | 1.73 | 0.71 |
| 1:A:47:GLU:N | 1:A:47:GLU:OE2 | 2.24 | 0.71 |
| 3:C:-157:ASP:HB3 | 3:C:-22:ARG:HE | 1.56 | 0.70 |
| 1:A:38:ARG:HH11 | 1:A:56:GLY:HA3 | 1.55 | 0.70 |
| 1:A:396:ILE:CG1 | 1:A:673:LEU:HD11 | 2.22 | 0.70 |
| 3:C:-95:ASP:O | 3:C:-91:ARG:HG3 | 1.92 | 0.70 |
| 1:A:674:LYS:O | 1:A:675:GLU:HG3 | 1.91 | 0.69 |
| 3:C:-50:SER:O | 3:C:-46:LEU:HD12 | 1.94 | 0.68 |
| 1:A:798:THR:HG23 | 1:A:800:GLU:H | 1.57 | 0.68 |
| 1:A:419:ARG:HG3 | 1:A:420:GLU:N | 2.09 | 0.66 |
| 1:A:81:THR:HG22 | 1:A:83:LYS:H | 1.60 | 0.66 |
| 1:A:986:ASP:OD1 | 1:A:989:ARG:NH1 | 2.27 | 0.66 |
| 2:B:1312:MET:HB2 | 2:B:1331:GLY:HA3 | 1.77 | 0.66 |
| 1:A:1061:VAL:HG13 | 1:A:1104:LYS:HD2 | 1.77 | 0.65 |
| 1:A:1057:ARG:HA | 1:A:1060:LYS:HE3 | 1.78 | 0.64 |
| 1:A:190:VAL:HG21 | 1:A:231:ILE:HD13 | 1.79 | 0.64 |
| 1:A:374:GLN:OE1 | 1:A:391:ARG:HB2 | 1.98 | 0.64 |
| 1:A:419:ARG:HG3 | 1:A:420:GLU:H | 1.64 | 0.63 |
| 3:C:-89:ILE:HG23 | 3:C:-83:LEU:HB3 | 1.78 | 0.63 |
| 2:B:1224:LYS:HE2 | 2:B:1260:MET:HG2 | 1.79 | 0.63 |
| 2:B:1226:ASN:HA | 2:B:1263:SER:HB2 | 1.80 | 0.63 |
| 1:A:891:TYR:CE1 | 1:A:901:THR:HG22 | 2.33 | 0.63 |
| 2:B:1262:ILE:HB | 2:B:1276:ASN:HB2 | 1.80 | 0.63 |
| 3:C:-75:ASP:N | 3:C:-75:ASP:OD1 | 2.31 | 0.63 |
| 1:A:413:LEU:HD11 | 1:A:468:LEU:HD22 | 1.81 | 0.62 |
| 1:A:339:ASP:OD2 | 6:A:1303:HOH:O | 2.16 | 0.62 |
| 2:B:1204:ASP:HB3 | 2:B:1207:THR:HG22 | 1.80 | 0.62 |
| 3:C:-23:ASN:O | 3:C:-19:ARG:HG3 | 2.01 | 0.61 |
| 1:A:498:ILE:HA | 1:A:512:VAL:HG12 | 1.82 | 0.61 |
| 1:A:578:HIS:NE2 | 1:A:580:GLU:HG2 | 2.15 | 0.60 |

*Continued on next page...*

CONFIDENTIAL VALIDATION REPORT

Continued from previous page...

| Atom-1 | Atom-2 | Interatomic distance (Å) | Clash overlap (Å) |
|---|---|---|---|
| 1:A:11:LYS:HZ1 | 1:A:38:ARG:HH21 | 1.50 | 0.60 |
| 1:A:672:ASN:O | 1:A:673:LEU:HD12 | 2.02 | 0.60 |
| 1:A:432:GLN:HB2 | 1:A:453:ASP:O | 2.02 | 0.60 |
| 3:C:-45:GLN:O | 3:C:-43:LYS:HG2 | 2.02 | 0.60 |
| 1:A:231:ILE:HD12 | 1:A:240:HIS:CD2 | 2.37 | 0.59 |
| 1:A:727:GLN:HG2 | 1:A:818:SER:OG | 2.02 | 0.59 |
| 2:B:1074:HIS:HD1 | 2:B:1345:ILE:HG12 | 1.67 | 0.58 |
| 1:A:389:ILE:HD13 | 1:A:713:ARG:HD2 | 1.85 | 0.58 |
| 2:B:1351:LYS:HD2 | 2:B:1351:LYS:N | 2.18 | 0.58 |
| 3:C:-151:LYS:HG2 | 3:C:-110:VAL:HG22 | 1.84 | 0.58 |
| 2:B:1047:PRO:HD2 | 2:B:1053:ARG:HG2 | 1.84 | 0.58 |
| 2:B:1393:ARG:CZ | 2:B:1398:LEU:HD21 | 2.31 | 0.57 |
| 1:A:285:LEU:HB3 | 1:A:297:LEU:CD1 | 2.32 | 0.57 |
| 1:A:491:LYS:HE2 | 1:A:491:LYS:CA | 2.29 | 0.57 |
| 1:A:589:ARG:NH1 | 1:A:637:VAL:HG22 | 2.20 | 0.57 |
| 1:A:641:PHE:HB3 | 1:A:681:PRO:HG3 | 1.86 | 0.57 |
| 1:A:885:ASN:O | 1:A:910:MET:HA | 2.05 | 0.56 |
| 2:B:1259:ASN:HD21 | 2:B:1296:GLN:NE2 | 2.02 | 0.56 |
| 2:B:1357:LEU:HD13 | 2:B:1366:LEU:HD11 | 1.86 | 0.56 |
| 1:A:396:ILE:HG12 | 1:A:673:LEU:HD11 | 1.86 | 0.56 |
| 1:A:485:ALA:O | 1:A:487:VAL:HG23 | 2.05 | 0.56 |
| 1:A:432:GLN:HG2 | 1:A:434:ARG:NH1 | 2.20 | 0.56 |
| 1:A:263:ARG:HG3 | 1:A:271:TYR:CE1 | 2.41 | 0.55 |
| 1:A:492:GLU:HG3 | 1:A:512:VAL:HG21 | 1.88 | 0.55 |
| 3:C:-45:GLN:C | 3:C:-43:LYS:H | 2.08 | 0.55 |
| 2:B:1081:ARG:HH22 | 2:B:1397:GLU:CD | 2.09 | 0.55 |
| 1:A:446:THR:HG22 | 1:A:447:GLU:H | 1.72 | 0.55 |
| 1:A:425:LEU:HD21 | 1:A:427:LEU:HD21 | 1.88 | 0.55 |
| 1:A:719:GLU:HG3 | 1:A:755:SER:HB2 | 1.87 | 0.54 |
| 2:B:1259:ASN:ND2 | 2:B:1296:GLN:HE21 | 2.05 | 0.54 |
| 2:B:1221:ASN:N | 2:B:1240:ASP:OD2 | 2.41 | 0.54 |
| 1:A:676:VAL:HG11 | 1:A:693:LEU:HD23 | 1.90 | 0.54 |
| 1:A:993:GLN:OE1 | 1:A:993:GLN:HA | 2.08 | 0.54 |
| 2:B:1185:SER:OG | 2:B:1188:SER:O | 2.21 | 0.54 |
| 1:A:448:LEU:HA | 1:A:484:LYS:NZ | 2.23 | 0.54 |
| 1:A:642:ARG:NH1 | 1:A:647:THR:HB | 2.22 | 0.54 |
| 1:A:745:THR:HG23 | 1:A:782:PHE:HE2 | 1.73 | 0.54 |
| 3:C:-77:SER:O | 3:C:-43:LYS:CE | 2.56 | 0.54 |
| 1:A:1113:GLN:OE1 | 1:A:1123:GLU:CA | 2.49 | 0.54 |
| 2:B:1225:ARG:HD2 | 2:B:1261:ASN:HB3 | 1.90 | 0.54 |
| 2:B:1236:LEU:HD13 | 2:B:1243:LEU:HD21 | 1.89 | 0.54 |

Continued on next page...

CONFIDENTIAL VALIDATION REPORT

Continued from previous page...

| Atom-1 | Atom-2 | Interatomic distance (Å) | Clash overlap (Å) |
|---|---|---|---|
| 1:A:1128:ASP:O | 1:A:1132:VAL:HG23 | 2.08 | 0.53 |
| 1:A:385:GLY:HA3 | 1:A:719:GLU:O | 2.08 | 0.53 |
| 1:A:561:TRP:HB3 | 1:A:562:THR:HG23 | 1.90 | 0.53 |
| 1:A:81:THR:HG22 | 1:A:83:LYS:N | 2.23 | 0.53 |
| 2:B:1081:ARG:NH2 | 2:B:1397:GLU:OE1 | 2.41 | 0.53 |
| 1:A:690:SER:OG | 1:A:691:LEU:N | 2.42 | 0.53 |
| 3:C:15:ARG:NH2 | 3:C:72:ASP:OD1 | 2.42 | 0.53 |
| 1:A:246:LEU:HD13 | 1:A:297:LEU:HD23 | 1.91 | 0.53 |
| 1:A:886:SER:O | 1:A:908:ASN:HB2 | 2.09 | 0.53 |
| 3:C:-12:THR:OG1 | 3:C:-10:THR:HG22 | 2.08 | 0.53 |
| 3:C:57:ASP:HB2 | 3:C:60:TRP:CD2 | 2.43 | 0.53 |
| 1:A:589:ARG:HH12 | 1:A:637:VAL:HG22 | 1.74 | 0.52 |
| 1:A:782:PHE:C | 1:A:782:PHE:CD1 | 2.83 | 0.52 |
| 1:A:328:LEU:HD22 | 1:A:381:ALA:HB2 | 1.91 | 0.52 |
| 2:B:1074:HIS:NE2 | 2:B:1343:LYS:HE3 | 2.25 | 0.52 |
| 3:C:-77:SER:O | 3:C:-43:LYS:HE2 | 2.09 | 0.51 |
| 1:A:381:ALA:O | 1:A:720:SER:OG | 2.22 | 0.51 |
| 1:A:38:ARG:HH11 | 1:A:56:GLY:CA | 2.22 | 0.51 |
| 1:A:474:ALA:HB2 | 6:A:1315:HOH:O | 2.08 | 0.51 |
| 1:A:492:GLU:HB3 | 6:A:1301:HOH:O | 2.10 | 0.51 |
| 1:A:127:GLU:HB3 | 1:A:129:ARG:HD3 | 1.93 | 0.51 |
| 1:A:421:THR:OG1 | 1:A:423:ASP:OD1 | 2.28 | 0.51 |
| 1:A:504:ASN:HD22 | 1:A:545:PRO:HD3 | 1.75 | 0.51 |
| 1:A:396:ILE:O | 1:A:396:ILE:HD12 | 2.11 | 0.51 |
| 1:A:148:ASP:N | 1:A:148:ASP:OD1 | 2.38 | 0.50 |
| 3:C:-161:MET:HG3 | 3:C:-6:TYR:CE2 | 2.46 | 0.50 |
| 1:A:188:ARG:NH1 | 1:A:216:ALA:O | 2.44 | 0.50 |
| 1:A:491:LYS:HD2 | 1:A:495:ALA:HA | 1.93 | 0.50 |
| 1:A:467:GLN:NE2 | 1:A:478:LEU:HD21 | 2.27 | 0.50 |
| 1:A:907:ASN:OD1 | 2:B:1391:ARG:NH2 | 2.44 | 0.50 |
| 2:B:1134:HIS:CG | 2:B:1154:ALA:HB2 | 2.47 | 0.50 |
| 3:C:-44:GLN:O | 3:C:-42:ARG:NE | 2.45 | 0.50 |
| 3:C:-96:VAL:O | 3:C:-92:VAL:HG12 | 2.12 | 0.50 |
| 1:A:356:LEU:HD21 | 1:A:712:ILE:HD13 | 1.94 | 0.50 |
| 1:A:538:VAL:HA | 1:A:560:LEU:HD23 | 1.94 | 0.50 |
| 1:A:507:GLN:NE2 | 1:A:552:LEU:HG | 2.26 | 0.50 |
| 1:A:72:GLU:OE2 | 1:A:103:ARG:NH2 | 2.37 | 0.49 |
| 1:A:231:ILE:HD12 | 1:A:240:HIS:HD2 | 1.76 | 0.49 |
| 1:A:467:GLN:HE21 | 1:A:478:LEU:HD21 | 1.77 | 0.49 |
| 1:A:453:ASP:OD1 | 1:A:453:ASP:N | 2.44 | 0.49 |
| 3:C:-108:THR:O | 3:C:-105:GLU:N | 2.43 | 0.49 |

Continued on next page...

CONFIDENTIAL VALIDATION REPORT

Continued from previous page...

| Atom-1 | Atom-2 | Interatomic distance (Å) | Clash overlap (Å) |
|---|---|---|---|
| 3:C:-40:ASP:O | 3:C:-36:VAL:HG23 | 2.12 | 0.49 |
| 1:A:55:VAL:HG21 | 1:A:1065:VAL:HG21 | 1.93 | 0.49 |
| 1:A:58:TYR:HB3 | 1:A:1073:TRP:HB2 | 1.95 | 0.49 |
| 2:B:1156:TRP:CD2 | 3:C:34:GLU:HG2 | 2.48 | 0.49 |
| 1:A:944:GLU:OE2 | 1:A:947:ARG:HD2 | 2.13 | 0.49 |
| 3:C:-21:ALA:O | 3:C:-17:ILE:HG13 | 2.13 | 0.49 |
| 1:A:318:ASP:OD1 | 4:A:1204:GOL:H31 | 2.12 | 0.49 |
| 1:A:492:GLU:CB | 6:A:1301:HOH:O | 2.60 | 0.49 |
| 2:B:1245:ASP:HB2 | 2:B:1252:ILE:HD11 | 1.93 | 0.49 |
| 3:C:-162:GLU:O | 3:C:-158:ILE:HD12 | 2.13 | 0.49 |
| 1:A:695:ASN:OD1 | 1:A:698:THR:N | 2.27 | 0.48 |
| 1:A:828:TYR:CD1 | 1:A:852:GLN:HB3 | 2.48 | 0.48 |
| 1:A:36:ASN:O | 1:A:60:LYS:HA | 2.14 | 0.48 |
| 2:B:1192:VAL:HG23 | 2:B:1205:ILE:HG22 | 1.94 | 0.48 |
| 1:A:749:THR:HG21 | 1:A:786:VAL:HG11 | 1.96 | 0.48 |
| 3:C:-62:GLN:HB2 | 3:C:-22:ARG:NH1 | 2.28 | 0.48 |
| 1:A:607:GLY:HA2 | 1:A:635:PRO:HB3 | 1.95 | 0.48 |
| 1:A:480:SER:O | 1:A:484:LYS:HA | 2.14 | 0.48 |
| 3:C:76:ARG:NE | 6:C:202:HOH:O | 2.42 | 0.48 |
| 1:A:1056:ASN:O | 1:A:1060:LYS:HG3 | 2.14 | 0.47 |
| 1:A:674:LYS:O | 1:A:675:GLU:CG | 2.59 | 0.47 |
| 1:A:1011:SER:OG | 1:A:1013:VAL:HG22 | 2.14 | 0.47 |
| 1:A:181:VAL:HG22 | 1:A:190:VAL:HG22 | 1.97 | 0.47 |
| 1:A:332:GLN:HB3 | 1:A:334:VAL:HG23 | 1.95 | 0.47 |
| 2:B:1224:LYS:HG3 | 2:B:1260:MET:O | 2.14 | 0.47 |
| 1:A:578:HIS:CD2 | 1:A:580:GLU:HG2 | 2.49 | 0.47 |
| 1:A:777:PRO:HG3 | 1:A:837:TYR:CD1 | 2.50 | 0.47 |
| 2:B:1352:ARG:HB3 | 2:B:1371:ASN:O | 2.15 | 0.47 |
| 1:A:143:ILE:HG12 | 1:A:154:ALA:HB2 | 1.97 | 0.47 |
| 2:B:1120:TYR:CZ | 2:B:1127:GLU:HG3 | 2.50 | 0.47 |
| 1:A:1097:PHE:O | 1:A:1100:ILE:HG12 | 2.15 | 0.47 |
| 1:A:368:GLU:HA | 1:A:368:GLU:OE1 | 2.14 | 0.47 |
| 2:B:1046:ALA:HB1 | 2:B:1071:PHE:HE2 | 1.80 | 0.47 |
| 2:B:1259:ASN:HD22 | 2:B:1276:ASN:ND2 | 2.12 | 0.47 |
| 2:B:1074:HIS:ND1 | 2:B:1345:ILE:HG12 | 2.28 | 0.47 |
| 3:C:-84:LYS:HB3 | 3:C:-84:LYS:HE3 | 1.65 | 0.47 |
| 1:A:1131:LYS:HD3 | 6:A:1302:HOH:O | 2.13 | 0.46 |
| 1:A:474:ALA:O | 1:A:475:SER:HB2 | 2.16 | 0.46 |
| 1:A:468:LEU:HD21 | 1:A:481:GLN:NE2 | 2.31 | 0.46 |
| 1:A:168:LYS:HE3 | 6:A:1399:HOH:O | 2.14 | 0.46 |
| 1:A:440:GLY:O | 1:A:686:GLY:HA3 | 2.16 | 0.46 |

Continued on next page...

CONFIDENTIAL VALIDATION REPORT

Continued from previous page...

| Atom-1 | Atom-2 | Interatomic distance (Å) | Clash overlap (Å) |
|---|---|---|---|
| 1:A:727:GLN:HG3 | 1:A:829:PHE:CZ | 2.51 | 0.46 |
| 1:A:828:TYR:HD1 | 1:A:852:GLN:HB3 | 1.79 | 0.46 |
| 1:A:906:TYR:CZ | 2:B:1396:GLU:OE1 | 2.62 | 0.46 |
| 3:C:-102:LYS:O | 3:C:-98:GLN:HG3 | 2.15 | 0.46 |
| 1:A:1136:LEU:O | 1:A:1139:ILE:HG12 | 2.15 | 0.46 |
| 1:A:190:VAL:HG21 | 1:A:231:ILE:CD1 | 2.45 | 0.46 |
| 1:A:40:GLU:HB3 | 1:A:42:TYR:CE1 | 2.51 | 0.46 |
| 1:A:490:TRP:CG | 1:A:519:LEU:HD21 | 2.50 | 0.46 |
| 1:A:770:LEU:HG | 1:A:865:GLU:HG3 | 1.98 | 0.46 |
| 1:A:417:PRO:HG3 | 1:A:481:GLN:OE1 | 2.16 | 0.46 |
| 1:A:560:LEU:HD12 | 1:A:567:ARG:HH11 | 1.81 | 0.46 |
| 3:C:-30:ARG:HH21 | 3:C:-26:GLN:HG2 | 1.81 | 0.46 |
| 1:A:131:ILE:HG13 | 1:A:145:LEU:HD11 | 1.98 | 0.45 |
| 1:A:255:GLN:HB3 | 1:A:279:ARG:NH2 | 2.30 | 0.45 |
| 3:C:-75:ASP:O | 3:C:-71:ARG:HG2 | 2.17 | 0.45 |
| 1:A:777:PRO:HG3 | 1:A:837:TYR:CE1 | 2.51 | 0.45 |
| 2:B:1224:LYS:HB2 | 2:B:1261:ASN:HA | 1.98 | 0.45 |
| 1:A:208:LYS:HD3 | 1:A:208:LYS:HA | 1.79 | 0.45 |
| 1:A:334:VAL:HG11 | 1:A:347:VAL:HG13 | 1.98 | 0.45 |
| 1:A:491:LYS:CD | 1:A:495:ALA:HA | 2.47 | 0.45 |
| 2:B:1074:HIS:HA | 2:B:1345:ILE:HG23 | 1.98 | 0.45 |
| 3:C:-51:ASN:O | 3:C:-47:MET:CG | 2.65 | 0.45 |
| 1:A:124:ILE:HG12 | 1:A:131:ILE:HG12 | 1.98 | 0.45 |
| 1:A:921:ILE:HB | 1:A:933:LEU:HB2 | 1.99 | 0.45 |
| 2:B:1341:ASP:OD2 | 2:B:1343:LYS:HE2 | 2.17 | 0.45 |
| 2:B:1105:GLU:OE1 | 2:B:1360:ASP:HB2 | 2.17 | 0.45 |
| 3:C:-12:THR:HG22 | 6:C:213:HOH:O | 2.17 | 0.45 |
| 1:A:560:LEU:CD1 | 1:A:567:ARG:HH11 | 2.30 | 0.45 |
| 1:A:637:VAL:HB | 1:A:652:CYS:HB2 | 1.99 | 0.45 |
| 1:A:857:LYS:NZ | 6:A:1304:HOH:O | 2.33 | 0.44 |
| 2:B:1119:LEU:CD2 | 3:C:4:VAL:HG21 | 2.47 | 0.44 |
| 1:A:1125:THR:HG23 | 1:A:1128:ASP:H | 1.81 | 0.44 |
| 1:A:439:ASN:OD1 | 1:A:439:ASN:N | 2.51 | 0.44 |
| 1:A:520:GLN:HG3 | 1:A:529:ILE:HG13 | 1.99 | 0.44 |
| 1:A:891:TYR:HE1 | 1:A:901:THR:HG22 | 1.79 | 0.44 |
| 2:B:1298:ARG:O | 2:B:1310:GLY:HA2 | 2.17 | 0.44 |
| 1:A:155:PHE:C | 1:A:155:PHE:CD1 | 2.91 | 0.44 |
| 1:A:597:GLU:OE2 | 1:A:661:SER:OG | 2.28 | 0.44 |
| 1:A:606:LEU:HD11 | 1:A:612:PHE:CE1 | 2.43 | 0.44 |
| 3:C:-45:GLN:C | 3:C:-43:LYS:N | 2.69 | 0.44 |
| 1:A:112:ILE:HD12 | 2:B:1232:PRO:HB2 | 2.00 | 0.44 |

Continued on next page...

CONFIDENTIAL VALIDATION REPORT

*Continued from previous page...*

| Atom-1 | Atom-2 | Interatomic distance (Å) | Clash overlap (Å) |
|---|---|---|---|
| 1:A:182:TYR:OH | 1:A:209:GLN:OE1 | 2.25 | 0.44 |
| 1:A:448:LEU:HA | 1:A:484:LYS:HZ1 | 1.82 | 0.44 |
| 2:B:1388:GLU:HG2 | 2:B:1391:ARG:HG3 | 1.99 | 0.44 |
| 3:C:-30:ARG:HH21 | 3:C:-26:GLN:CG | 2.30 | 0.44 |
| 1:A:851:PHE:HB3 | 1:A:858:LEU:HD11 | 2.00 | 0.44 |
| 1:A:586:ILE:HD12 | 1:A:587:ILE:H | 1.82 | 0.43 |
| 2:B:1076:ILE:O | 2:B:1390:GLY:HA2 | 2.17 | 0.43 |
| 2:B:1146:ASP:OD1 | 2:B:1146:ASP:N | 2.49 | 0.43 |
| 2:B:1332:SER:HB3 | 2:B:1375:MET:SD | 2.58 | 0.43 |
| 1:A:23:PHE:CE2 | 1:A:91:TYR:HB2 | 2.53 | 0.43 |
| 1:A:477:ARG:NH2 | 1:A:486:LEU:HD22 | 2.32 | 0.43 |
| 1:A:1030:PHE:CD1 | 1:A:1030:PHE:C | 2.92 | 0.43 |
| 1:A:1106:GLN:O | 1:A:1109:VAL:HG12 | 2.18 | 0.43 |
| 1:A:158:ARG:HD3 | 2:B:1283:ARG:HG2 | 1.99 | 0.43 |
| 3:C:-140:ILE:HG13 | 3:C:-139:GLY:N | 2.33 | 0.43 |
| 3:C:-140:ILE:HG21 | 3:C:-121:LEU:HD13 | 2.00 | 0.43 |
| 2:B:1236:LEU:HB3 | 2:B:1243:LEU:HD21 | 2.00 | 0.43 |
| 3:C:-117:ILE:HD13 | 3:C:-105:GLU:HB3 | 2.00 | 0.43 |
| 1:A:334:VAL:CG1 | 1:A:347:VAL:HG13 | 2.48 | 0.43 |
| 1:A:432:GLN:HG2 | 1:A:434:ARG:HH12 | 1.84 | 0.43 |
| 1:A:605:ALA:HB2 | 1:A:611:LEU:HD23 | 2.01 | 0.43 |
| 1:A:811:GLU:OE2 | 1:A:847:ARG:HD3 | 2.18 | 0.43 |
| 2:B:1109:MET:HG2 | 2:B:1119:LEU:HG | 2.01 | 0.43 |
| 3:C:-162:GLU:OE1 | 3:C:-159:ARG:NE | 2.48 | 0.43 |
| 1:A:610:ALA:HB1 | 1:A:628:LYS:HE3 | 2.00 | 0.43 |
| 2:B:1190:ASP:O | 2:B:1205:ILE:HG12 | 2.19 | 0.43 |
| 2:B:1231:ASN:ND2 | 2:B:1233:THR:OG1 | 2.49 | 0.43 |
| 1:A:353:PHE:CG | 4:A:1203:GOL:H11 | 2.54 | 0.43 |
| 1:A:414:ARG:NH2 | 1:A:419:ARG:O | 2.49 | 0.43 |
| 3:C:4:VAL:HG22 | 3:C:5:PRO:HD2 | 2.01 | 0.43 |
| 1:A:554:PRO:O | 1:A:570:LYS:HD2 | 2.19 | 0.42 |
| 2:B:1081:ARG:HH21 | 2:B:1391:ARG:NH1 | 2.18 | 0.42 |
| 2:B:1291:VAL:HG23 | 2:B:1291:VAL:O | 2.19 | 0.42 |
| 1:A:261:HIS:HA | 1:A:272:LEU:O | 2.20 | 0.42 |
| 1:A:745:THR:HG23 | 1:A:782:PHE:CE2 | 2.54 | 0.42 |
| 1:A:969:GLU:OE2 | 1:A:973:ASN:HB2 | 2.19 | 0.42 |
| 2:B:1151:LEU:HB3 | 2:B:1182:VAL:HG13 | 2.01 | 0.42 |
| 1:A:359:ILE:HG21 | 1:A:362:MET:CE | 2.49 | 0.42 |
| 1:A:358:PRO:O | 1:A:379:SER:HA | 2.20 | 0.42 |
| 3:C:-42:ARG:HA | 3:C:-42:ARG:HD3 | 1.82 | 0.42 |
| 1:A:425:LEU:HD12 | 1:A:682:LEU:HD12 | 2.00 | 0.42 |

*Continued on next page...*

CONFIDENTIAL VALIDATION REPORT

*Continued from previous page...*

| Atom-1 | Atom-2 | Interatomic distance (Å) | Clash overlap (Å) |
|---|---|---|---|
| 1:A:478:LEU:HD12 | 1:A:479:VAL:H | 1.84 | 0.42 |
| 2:B:1214:LEU:HD22 | 2:B:1244:TRP:CE3 | 2.54 | 0.42 |
| 3:C:-161:MET:HE2 | 3:C:-6:TYR:OH | 2.20 | 0.42 |
| 1:A:147:ARG:O | 1:A:150:LYS:NZ | 2.53 | 0.42 |
| 1:A:654:ASP:OD1 | 1:A:654:ASP:N | 2.53 | 0.42 |
| 1:A:996:GLY:O | 1:A:997:LEU:HD23 | 2.20 | 0.42 |
| 1:A:609:GLY:HA3 | 1:A:632:GLY:O | 2.20 | 0.42 |
| 2:B:1109:MET:HE1 | 2:B:1150:LEU:HD22 | 2.01 | 0.42 |
| 2:B:1204:ASP:HB3 | 2:B:1207:THR:CG2 | 2.48 | 0.42 |
| 2:B:1396:GLU:HG2 | 6:B:1626:HOH:O | 2.20 | 0.42 |
| 3:C:-140:ILE:HB | 3:C:-134:LEU:HD11 | 2.01 | 0.42 |
| 1:A:133:LEU:HB2 | 1:A:141:LYS:HB3 | 2.01 | 0.42 |
| 1:A:181:VAL:HA | 1:A:189:HIS:O | 2.20 | 0.42 |
| 1:A:413:LEU:HD13 | 1:A:461:GLY:HA2 | 2.02 | 0.41 |
| 1:A:491:LYS:HD3 | 1:A:492:GLU:N | 2.35 | 0.41 |
| 3:C:14:SER:CB | 3:C:17:VAL:HG12 | 2.45 | 0.41 |
| 1:A:396:ILE:HG12 | 1:A:673:LEU:HD21 | 2.02 | 0.41 |
| 1:A:650:PHE:HD1 | 1:A:651:ALA:N | 2.19 | 0.41 |
| 1:A:288:GLU:HG3 | 1:A:298:LYS:HB2 | 2.02 | 0.41 |
| 1:A:446:THR:HG22 | 1:A:447:GLU:N | 2.35 | 0.41 |
| 1:A:654:ASP:HA | 1:A:675:GLU:HG2 | 2.01 | 0.41 |
| 1:A:719:GLU:HG2 | 1:A:739:ARG:HB3 | 2.01 | 0.41 |
| 1:A:719:GLU:OE2 | 1:A:757:SER:OG | 2.26 | 0.41 |
| 1:A:394:ILE:HG12 | 1:A:708:GLN:HA | 2.02 | 0.41 |
| 2:B:1067:ASP:N | 2:B:1067:ASP:OD1 | 2.54 | 0.41 |
| 1:A:14:ALA:HB1 | 1:A:327:ARG:HB2 | 2.03 | 0.41 |
| 1:A:40:GLU:HB3 | 1:A:42:TYR:HE1 | 1.85 | 0.41 |
| 1:A:451:PHE:HA | 1:A:470:GLN:OE1 | 2.21 | 0.41 |
| 1:A:512:VAL:HG23 | 1:A:512:VAL:O | 2.21 | 0.41 |
| 3:C:14:SER:HB3 | 3:C:17:VAL:CG1 | 2.46 | 0.41 |
| 1:A:10:GLN:HB3 | 1:A:1037:ILE:HB | 2.03 | 0.41 |
| 1:A:112:ILE:HG23 | 1:A:112:ILE:O | 2.21 | 0.41 |
| 1:A:743:GLN:HG2 | 1:A:782:PHE:HA | 2.03 | 0.41 |
| 1:A:889:ARG:HD2 | 1:A:891:TYR:OH | 2.21 | 0.41 |
| 2:B:1109:MET:HE3 | 2:B:1117:LEU:HD21 | 2.02 | 0.41 |
| 2:B:1178:GLU:OE1 | 2:B:1196:LYS:NZ | 2.54 | 0.41 |
| 2:B:1333:SER:OG | 2:B:1349:ASP:HA | 2.21 | 0.41 |
| 1:A:650:PHE:CD2 | 1:A:679:MET:HG3 | 2.56 | 0.41 |
| 1:A:927:MET:HG3 | 1:A:953:TRP:CE2 | 2.56 | 0.41 |
| 3:C:-45:GLN:O | 3:C:-43:LYS:N | 2.53 | 0.41 |
| 3:C:31:LEU:HD21 | 3:C:77:THR:HB | 2.02 | 0.40 |

*Continued on next page...*

CONFIDENTIAL VALIDATION REPORT

*Continued from previous page...*

| Atom-1 | Atom-2 | Interatomic distance (Å) | Clash overlap (Å) |
|---|---|---|---|
| 1:A:58:TYR:HB3 | 1:A:1073:TRP:CB | 2.50 | 0.40 |
| 1:A:284:LEU:HD13 | 1:A:301[A]:ARG:NH2 | 2.36 | 0.40 |
| 2:B:1104:ARG:C | 2:B:1105:GLU:HG2 | 2.41 | 0.40 |
| 1:A:983:ALA:HB1 | 1:A:988:GLU:HB3 | 2.04 | 0.40 |
| 2:B:1104:ARG:HD3 | 3:C:2:GLU:HB2 | 2.02 | 0.40 |

There are no symmetry-related clashes.

#### 5.3 Torsion angles (i)

##### 5.3.1 Protein backbone (i)

In the following table, the Percentiles column shows the percent Ramachandran outliers of the chain as a percentile score with respect to all X-ray entries followed by that with respect to entries of similar resolution.

The Analysed column shows the number of residues for which the backbone conformation was analysed, and the total number of residues.

| Mol | Chain | Analysed | Favoured | Allowed | Outliers | Percentiles | |
|---|---|---|---|---|---|---|---|
| 1 | A | 1117/1142 (98%) | 1061 (95%) | 56 (5%) | 0 | 100 | 100 |
| 2 | B | 329/360 (91%) | 309 (94%) | 20 (6%) | 0 | 100 | 100 |
| 3 | C | 255/258 (99%) | 247 (97%) | 7 (3%) | 1 (0%) | 34 | 53 |
| All | All | 1701/1760 (97%) | 1617 (95%) | 83 (5%) | 1 (0%) | 51 | 71 |

All (1) Ramachandran outliers are listed below:

| Mol | Chain | Res | Type |
|---|---|---|---|
| 3 | C | -44 | GLN |

##### 5.3.2 Protein sidechains (i)

In the following table, the Percentiles column shows the percent sidechain outliers of the chain as a percentile score with respect to all X-ray entries followed by that with respect to entries of similar resolution.

The Analysed column shows the number of residues for which the sidechain conformation was analysed, and the total number of residues.

CONFIDENTIAL VALIDATION REPORT

| Mol | Chain | Analysed | Rotameric | Outliers | Percentiles | |
|---|---|---|---|---|---|---|
| 1 | A | 987/1000 (99%) | 970 (98%) | 17 (2%) | 60 | 81 |
| 2 | B | 294/315 (93%) | 289 (98%) | 5 (2%) | 60 | 81 |
| 3 | C | 217/218 (100%) | 207 (95%) | 10 (5%) | 27 | 47 |
| All | All | 1498/1533 (98%) | 1466 (98%) | 32 (2%) | 53 | 76 |

All (32) residues with a non-rotameric sidechain are listed below:

| Mol | Chain | Res | Type |
|---|---|---|---|
| 1 | A | 5 | TYR |
| 1 | A | 111 | ARG |
| 1 | A | 155 | PHE |
| 1 | A | 204 | LYS |
| 1 | A | 287 | LYS |
| 1 | A | 335 | LYS |
| 1 | A | 366 | ASP |
| 1 | A | 423 | ASP |
| 1 | A | 491 | LYS |
| 1 | A | 650 | PHE |
| 1 | A | 680 | CYS |
| 1 | A | 699 | LEU |
| 1 | A | 713 | ARG |
| 1 | A | 782 | PHE |
| 1 | A | 857 | LYS |
| 1 | A | 1030 | PHE |
| 1 | A | 1123 | GLU |
| 2 | B | 1070 | CYS |
| 2 | B | 1167 | LYS |
| 2 | B | 1212 | LEU |
| 2 | B | 1283 | ARG |
| 2 | B | 1327 | LYS |
| 3 | C | -119 | LYS |
| 3 | C | -48 | ARG |
| 3 | C | -47 | MET |
| 3 | C | -46 | LEU |
| 3 | C | -43 | LYS |
| 3 | C | -32 | LYS |
| 3 | C | 9 | ARG |
| 3 | C | 41 | ARG |
| 3 | C | 57 | ASP |
| 3 | C | 89 | CYS |

Some sidechains can be flipped to improve hydrogen bonding and reduce clashes. All (3) such

CONFIDENTIAL VALIDATION REPORT

sidechains are listed below:

| Mol | Chain | Res | Type |
|---|---|---|---|
| 1 | A | 467 | GLN |
| 2 | B | 1296 | GLN |
| 3 | C | -45 | GLN |

##### 5.3.3 RNA (i)

There are no RNA molecules in this entry.

#### 5.4 Non-standard residues in protein, DNA, RNA chains (i)

There are no non-standard protein/DNA/RNA residues in this entry.

#### 5.5 Carbohydrates (i)

There are no monosaccharides in this entry.

#### 5.6 Ligand geometry (i)

Of 9 ligands modelled in this entry, 1 is monoatomic - leaving 8 for Mogul analysis.

In the following table, the Counts columns list the number of bonds (or angles) for which Mogul statistics could be retrieved, the number of bonds (or angles) that are observed in the model and the number of bonds (or angles) that are defined in the Chemical Component Dictionary. The Link column lists molecule types, if any, to which the group is linked. The Z score for a bond length (or angle) is the number of standard deviations the observed value is removed from the expected value. A bond length (or angle) with $|Z| > 2$ is considered an outlier worth inspection. RMSZ is the root-mean-square of all Z scores of the bond lengths (or angles).

| Mol | Type | Chain | Res | Link | Bond lengths | | | Bond angles | | |
|---|---|---|---|---|---|---|---|---|---|---|
| | | | | | Counts | RMSZ | $\#\|Z\| > 2$ | Counts | RMSZ | $\#\|Z\| > 2$ |
| 4 | GOL | A | 1206 | - | 5,5,5 | 0.92 | 0 | 5,5,5 | 0.97 | 0 |
| 4 | GOL | A | 1203 | - | 5,5,5 | 0.89 | 0 | 5,5,5 | 1.03 | 0 |
| 4 | GOL | C | 102 | - | 5,5,5 | 0.92 | 0 | 5,5,5 | 1.00 | 0 |
| 4 | GOL | B | 1501 | - | 5,5,5 | 0.90 | 0 | 5,5,5 | 1.00 | 0 |
| 4 | GOL | A | 1204 | - | 5,5,5 | 1.00 | 0 | 5,5,5 | 0.82 | 0 |
| 4 | GOL | A | 1201 | - | 5,5,5 | 0.90 | 0 | 5,5,5 | 1.01 | 0 |
| 4 | GOL | A | 1202 | - | 5,5,5 | 0.95 | 0 | 5,5,5 | 0.90 | 0 |
| 4 | GOL | A | 1205 | - | 5,5,5 | 0.93 | 0 | 5,5,5 | 0.97 | 0 |

In the following table, the Chirals column lists the number of chiral outliers, the number of chiral

CONFIDENTIAL VALIDATION REPORT

centers analysed, the number of these observed in the model and the number defined in the Chemical Component Dictionary. Similar counts are reported in the Torsion and Rings columns. '-' means no outliers of that kind were identified.

| Mol | Type | Chain | Res | Link | Chirals | Torsions | Rings |
|---|---|---|---|---|---|---|---|
| 4 | GOL | A | 1206 | - | - | 3/4/4/4 | - |
| 4 | GOL | A | 1203 | - | - | 4/4/4/4 | - |
| 4 | GOL | C | 102 | - | - | 0/4/4/4 | - |
| 4 | GOL | B | 1501 | - | - | 2/4/4/4 | - |
| 4 | GOL | A | 1204 | - | - | 3/4/4/4 | - |
| 4 | GOL | A | 1201 | - | - | 0/4/4/4 | - |
| 4 | GOL | A | 1202 | - | - | 4/4/4/4 | - |
| 4 | GOL | A | 1205 | - | - | 2/4/4/4 | - |

There are no bond length outliers.

There are no bond angle outliers.

There are no chirality outliers.

All (18) torsion outliers are listed below:

| Mol | Chain | Res | Type | Atoms |
|---|---|---|---|---|
| 4 | A | 1203 | GOL | C1-C2-C3-O3 |
| 4 | A | 1203 | GOL | O2-C2-C3-O3 |
| 4 | B | 1501 | GOL | O1-C1-C2-C3 |
| 4 | A | 1202 | GOL | O1-C1-C2-C3 |
| 4 | A | 1202 | GOL | C1-C2-C3-O3 |
| 4 | A | 1204 | GOL | O2-C2-C3-O3 |
| 4 | A | 1203 | GOL | O1-C1-C2-C3 |
| 4 | A | 1204 | GOL | O1-C1-C2-C3 |
| 4 | A | 1204 | GOL | C1-C2-C3-O3 |
| 4 | A | 1205 | GOL | O1-C1-C2-C3 |
| 4 | B | 1501 | GOL | O1-C1-C2-O2 |
| 4 | A | 1202 | GOL | O1-C1-C2-O2 |
| 4 | A | 1202 | GOL | O2-C2-C3-O3 |
| 4 | A | 1205 | GOL | O1-C1-C2-O2 |
| 4 | A | 1206 | GOL | O2-C2-C3-O3 |
| 4 | A | 1206 | GOL | O1-C1-C2-C3 |
| 4 | A | 1203 | GOL | O1-C1-C2-O2 |
| 4 | A | 1206 | GOL | C1-C2-C3-O3 |

There are no ring outliers.

2 monomers are involved in 2 short contacts:

CONFIDENTIAL VALIDATION REPORT

| Mol | Chain | Res | Type | Clashes | Symm-Clashes |
|---|---|---|---|---|---|
| 4 | A | 1203 | GOL | 1 | 0 |
| 4 | A | 1204 | GOL | 1 | 0 |

#### 5.7 Other polymers (i)

There are no such residues in this entry.

#### 5.8 Polymer linkage issues (i)

There are no chain breaks in this entry.

CONFIDENTIAL VALIDATION REPORT

### 6 Fit of model and data (i)

#### 6.1 Protein, DNA and RNA chains (i)

In the following table, the column labelled '#RSRZ> 2' contains the number (and percentage) of RSRZ outliers, followed by percent RSRZ outliers for the chain as percentile scores relative to all X-ray entries and entries of similar resolution. The OWAB column contains the minimum, median, $95^{th}$ percentile and maximum values of the occupancy-weighted average B-factor per residue. The column labelled 'Q< 0.9' lists the number of (and percentage) of residues with an average occupancy less than 0.9.

| Mol | Chain | Analysed | <RSRZ> | #RSRZ>2 | | | OWAB(Å$^2$) | Q<0.9 |
|---|---|---|---|---|---|---|---|---|
| 1 | A | 1124/1142 (98%) | 0.32 | 38 (3%) | 45 | 49 | 29, 61, 121, 175 | 0 |
| 2 | B | 335/360 (93%) | 0.27 | 7 (2%) | 63 | 67 | 39, 67, 99, 132 | 0 |
| 3 | C | 257/258 (99%) | 0.48 | 13 (5%) | 28 | 30 | 47, 86, 126, 149 | 0 |
| All | All | 1716/1760 (97%) | 0.33 | 58 (3%) | 45 | 49 | 29, 67, 121, 175 | 0 |

All (58) RSRZ outliers are listed below:

| Mol | Chain | Res | Type | RSRZ |
|---|---|---|---|---|
| 3 | C | 16 | VAL | 7.7 |
| 1 | A | 448 | LEU | 5.7 |
| 1 | A | 111 | ARG | 5.5 |
| 2 | B | 1398 | LEU | 4.9 |
| 1 | A | 644 | LEU | 4.8 |
| 1 | A | 438 | LEU | 4.0 |
| 1 | A | 419 | ARG | 3.9 |
| 1 | A | 775 | THR | 3.7 |
| 2 | B | 1376 | ASP | 3.7 |
| 1 | A | 549 | SER | 3.6 |
| 1 | A | 1124 | ALA | 3.6 |
| 1 | A | 685 | ASP | 3.6 |
| 2 | B | 1399 | ALA | 3.5 |
| 1 | A | 295 | VAL | 3.4 |
| 1 | A | 699 | LEU | 3.4 |
| 1 | A | 773 | SER | 3.3 |
| 1 | A | 435 | VAL | 3.2 |
| 3 | C | -36 | VAL | 3.1 |
| 1 | A | 402 | ILE | 3.0 |
| 1 | A | 381 | ALA | 3.0 |
| 3 | C | -83 | LEU | 3.0 |
| 1 | A | 413 | LEU | 2.9 |
| 1 | A | 427 | LEU | 2.9 |

*Continued on next page...*

CONFIDENTIAL VALIDATION REPORT

*Continued from previous page...*

| Mol | Chain | Res | Type | RSRZ |
|---|---|---|---|---|
| 1 | A | 686 | GLY | 2.9 |
| 3 | C | -163 | PHE | 2.9 |
| 1 | A | 548 | ASP | 2.9 |
| 1 | A | 526 | LEU | 2.8 |
| 1 | A | 439 | ASN | 2.8 |
| 1 | A | 691 | LEU | 2.7 |
| 1 | A | 437 | MET | 2.7 |
| 1 | A | 444 | GLU | 2.7 |
| 1 | A | 441 | GLU | 2.6 |
| 3 | C | -13 | ARG | 2.6 |
| 1 | A | 410 | LEU | 2.6 |
| 3 | C | -48 | ARG | 2.5 |
| 1 | A | 112 | ILE | 2.5 |
| 3 | C | -30 | ARG | 2.5 |
| 1 | A | 1014 | MET | 2.4 |
| 3 | C | -8 | ASP | 2.4 |
| 1 | A | 301[A] | ARG | 2.4 |
| 2 | B | 1375 | MET | 2.3 |
| 3 | C | 18 | PRO | 2.3 |
| 3 | C | -42 | ARG | 2.3 |
| 1 | A | 403 | ASP | 2.3 |
| 1 | A | 1114 | TYR | 2.3 |
| 2 | B | 1214 | LEU | 2.2 |
| 3 | C | -76 | LEU | 2.2 |
| 1 | A | 655 | ARG | 2.2 |
| 1 | A | 687 | TYR | 2.2 |
| 1 | A | 774 | SER | 2.2 |
| 3 | C | -74 | ALA | 2.1 |
| 1 | A | 404 | LEU | 2.1 |
| 3 | C | -89 | ILE | 2.1 |
| 2 | B | 1288 | LEU | 2.1 |
| 1 | A | 96 | GLU | 2.1 |
| 2 | B | 1327 | LYS | 2.1 |
| 1 | A | 469 | ILE | 2.1 |
| 1 | A | 101 | ILE | 2.0 |

#### 6.2 Non-standard residues in protein, DNA, RNA chains (i)

There are no non-standard protein/DNA/RNA residues in this entry.

CONFIDENTIAL VALIDATION REPORT

#### 6.3 Carbohydrates (i)

There are no monosaccharides in this entry.

#### 6.4 Ligands (i)

In the following table, the Atoms column lists the number of modelled atoms in the group and the number defined in the chemical component dictionary. The B-factors column lists the minimum, median, $95^{th}$ percentile and maximum values of B factors of atoms in the group. The column labelled 'Q< 0.9' lists the number of atoms with occupancy less than 0.9.

| Mol | Type | Chain | Res | Atoms | RSCC | RSR | B-factors(Å$^2$) | Q<0.9 |
|---|---|---|---|---|---|---|---|---|
| 4 | GOL | B | 1501 | 6/6 | 0.74 | 0.19 | 73,89,90,90 | 0 |
| 4 | GOL | A | 1204 | 6/6 | 0.79 | 0.24 | 69,73,79,81 | 0 |
| 4 | GOL | A | 1206 | 6/6 | 0.86 | 0.21 | 80,89,91,92 | 0 |
| 4 | GOL | C | 102 | 6/6 | 0.92 | 0.11 | 88,89,89,89 | 0 |
| 4 | GOL | A | 1205 | 6/6 | 0.92 | 0.18 | 69,73,74,76 | 0 |
| 4 | GOL | A | 1203 | 6/6 | 0.93 | 0.18 | 61,65,74,75 | 0 |
| 4 | GOL | A | 1201 | 6/6 | 0.94 | 0.25 | 44,58,66,77 | 0 |
| 5 | ZN | C | 101 | 1/1 | 0.96 | 0.17 | 79,79,79,79 | 0 |
| 4 | GOL | A | 1202 | 6/6 | 0.97 | 0.11 | 51,57,58,62 | 0 |

The following is a graphical depiction of the model fit to experimental electron density of all instances of the Ligand of Interest. In addition, ligands with molecular weight > 250 and outliers as shown on the geometry validation Tables will also be included. Each fit is shown from different orientation to approximate a three-dimensional view.

CONFIDENTIAL VALIDATION REPORT

**Electron density around ZN C 101:**

$2mF_o$-$DF_c$ (at 0.7 rmsd) in gray
$mF_o$-$DF_c$ (at 3 rmsd) in purple (negative)
and green (positive)

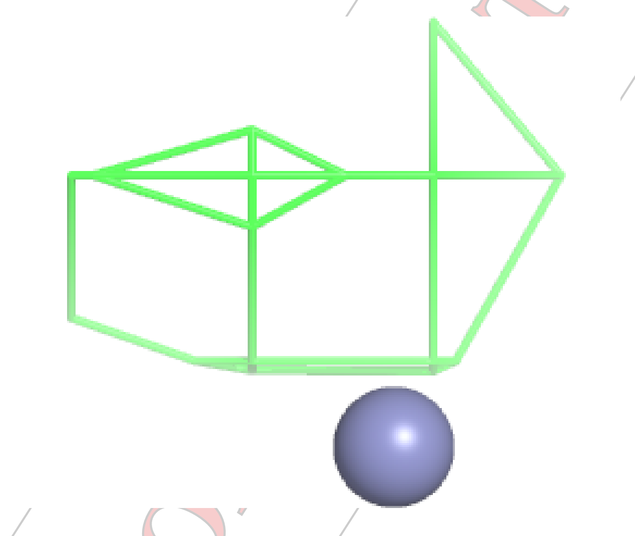

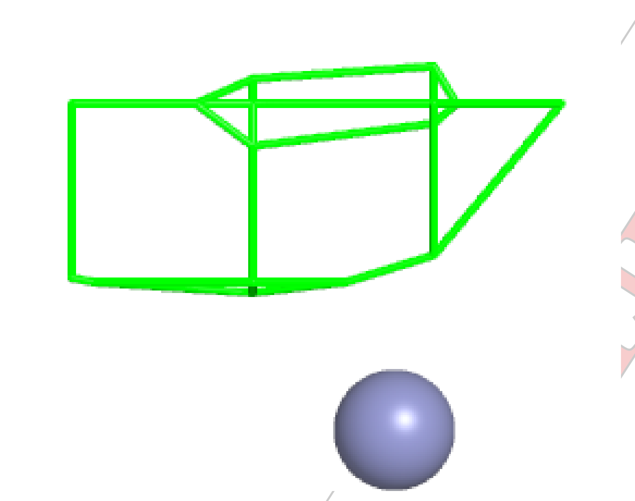

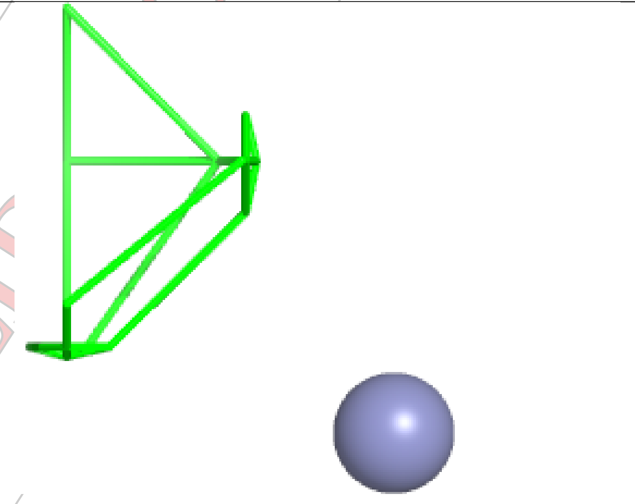

#### 6.5 Other polymers (i)

There are no such residues in this entry.

CONFIDENTIAL
